# Quantitative species delimitation, tests for gene flow, and migration models uncover complex, recent speciation in tree squirrels

**DOI:** 10.1101/2024.01.04.574244

**Authors:** Edson F. Abreu, Joyce R. Prado, Jesús E. Maldonado, Don E. Wilson, Alexandre R. Percequillo, Silvia E. Pavan

**Affiliations:** Department of Biology, Angelo State University, San Angelo, TX, USA; Department of Mammalogy, American Museum of Natural History, New York, NY, USA; Museu de Zoologia da Universidade de São Paulo, São Paulo, SP, Brazil; Center for Conservation Genomics, Smithsonian National Zoo and Conservation Biology Institute, Washington, DC, USA; Division of Mammals, National Museum of Natural History, Smithsonian Institution, Washington, DC, USA; Laboratório de Mamíferos, Departamento de Ciências Biológicas, Escola Superior de Agricultura Luiz de Queiroz, Universidade de São Paulo, Piracicaba, SP, Brazil; Department of Biological Sciences, California State Polytechnic University, Humboldt, Arcata, CA, USA

**Keywords:** Amazonia, diversification, mito-nuclear discordance, population genetics, Sciuridae, Ultraconserved Elements

## Abstract

Accurate estimates of species diversity are essential for all biodiversity research. Delimiting species and understanding the underlaying processes of speciation are also central components of systematic biology that outline our comprehension of the evolutionary mechanisms generating biodiversity. We obtained genomic data (Ultraconserved Elements and single nucleotide polymorphisms) for a widespread genus of South American tree squirrels (genus *Guerlinguetus*) to explore alternative hypotheses on species limits and to clarify recent and rapid speciation on continental-scale and dynamically evolving landscapes. Using a multilayered genomic approach that integrates fine-scale population genetic analyses with quantitative molecular species delimitation methods, we observed that (i) the most likely number of species within *Guerlinguetus* is six, contrasting with both classic morphological revision and mitochondrial species delimitation; (ii) incongruencies in species relationships still persist, which might be a response to population migration and gene flow taking place in lowlands of eastern Amazonia; and (iii) effective migration surfaces detected important geographic barriers associated with the major Amazonian riverine systems and the mountain ranges of the Guiana shield. In conclusion, we uncovered unexpected species diversity of keystone mammals that are critical in tree-seed predation and dispersal in one of the most fragile and threated ecosystems of the world, the tropical rainforests of South America. Our results corroborate recent findings suggesting that much of the extant species-level diversity in Amazonia is young, dating back to the Quaternary, while also reinforcing long-established hypotheses on the role of rivers and climate-driven forest dynamics triggering Amazonian speciation.

## 1. Introduction

Accurate estimates of species diversity lay the foundation and provide cascading effects in all biodiversity research (Stanton et al., 2019). Delimiting species and understanding the underlaying processes of speciation are also central components of systematic biology that outline our comprehension of the evolutionary mechanisms generating biodiversity (Mayr, 1968). However, neither of these tasks are trivial. Species delimitation concepts and methods changed vastly throughout centuries, from pre-evolutionary typological to genomic coalescent approaches, generating a lot of noise in evolutionary, ecological, and conservation practices (as reviewed by Stanton et al., 2019).

As genomic data proliferated for non-model organisms, they increasingly exposed cases of complex speciation (Chan et al., 2017; Giarla & Esselstyn, 2015; O’Connell et al., 2021). In such cases, gene flow resulting from continued interbreeding after initial divergence is widely recognized as a main element of speciation, producing, among other outcomes, intricate genomic evolution reflected in mito-nuclear discordances (Andersen et al., 2021; DeRaad et al., 2022). Unfortunately, until very recently most species delimitation approaches disregarded or were not able to account for the impacts of these interrelated phenomena. Species delimitation was (still is and will continue to be, in several cases) essentially performed by using specific elements of geographic groups as criteria to assess lineage independence, such as phenotypic distinctiveness, molecular divergence, and phylogenetic placement (Chan et al., 2017; D’Elía et al., 2019). These elements can be useful in cases where there are strong prezygotic barriers or when enough time has passed to allow the accumulation of fixed character differences. However, in many other cases, species delimitations can be misled when disregarding gene flow, which is currently seen as a pervasive, nearly inevitable component of speciation (Andersen et al., 2021; Leaché et al., 2019; Long & Kubatko, 2018).

Additionally, delimiting species that diversified rapidly and recently increases the challenge of recognizing clear boundaries and estimating the sequence of diversification events among lineages. Specifically, in recently diversified species, several regions of the genome might not have had enough time to evolve robust phylogenetic signals (Revell et al., 2008). If the diversification was also rapid, i.e., with short intervals between speciation events, this problem becomes even more intricate, as the most probable gene trees may confound the species branching history (Giarla & Esselstyn, 2015). Speciation can occur more rapidly in new, recently formed, and/or dynamic environments, where selection and pressure for adaptation are severe (Pereyra et al., 2009). Selection for local adaptations may act directly on loci underlying reproductive isolation or indirectly through the establishment of a genetic barrier to gene flow (Gavrilets, 2003). Furthermore, speciation is more likely to be triggered in dramatically changing environments, where genetic isolation occurs very quickly, driven by recently established barriers to migration (e.g., geographic or ecological).

The relatively young and dynamic formation of South American tropical rainforests, particularly in Amazonia, offered exceptional conditions for intense and rapid speciation events across the Tree of Life. This is evidenced by the strikingly high species richness in several groups of Amazonian organisms (Andrade-Silva et al., 2022; Cardoso et al., 2017; Mannion et al., 2014). The evolution of the Amazonas River basin, as seen from the perspective of processes generating local biodiversity, can be divided into two major temporal periods (Hoorn et al., 2010). The first, estimated to have occurred during the late Miocene, is characterized by the final stages of the Andes uplifting and the simultaneous formation of the main riverine systems, including the Amazonas River (Hoorn et al., 2010). The second period was driven by climatic fluctuations during the Pleistocene resulting in forest expansions and contractions that deeply affected Amazonian speciation history (Hoorn et al., 2010). However, recent findings are showing that the processes shaping the current fluvial organization may have permeated along the Plio-Pleistocene with, for instance, the transition from the lacustrine system in western Amazonia overlapping with Pleistocene glacial cycles (Cracraft et al., 2020; Ribas et al., 2012).

The Amazon Forest currently is separated geographically from the second largest tropical rainforest in South America, the Atlantic Forest, by a “dry diagonal” of open vegetation, which acts as an important barrier for biotic exchange between these two forested ecosystems (Raven & Axelrod, 1974). However, different lines of evidence show that historically the two rainforests were connected by continuous forest corridors that served as connecting routes for the dispersal of various groups (Batalha-Filho et al., 2013; Costa, 2003). Even so, information such as the period, frequency, and climatic conditions of these connections remain controversial (Ledo & Colli, 2017). The Atlantic Forest also harbors an impressive diversity of organisms (Myers et al., 2000), and shares with Amazonia a complex history of formation of species, with Pleistocene glacial cycles cited as responsible for the establishment of forest refugia (either in the mainland or in the continental shelf), that likely shaped the diversification in many taxa (Carnaval et al., 2009; Leite et al., 2016).

Tree squirrels of the genus *Guerlinguetus* represent a recent South American radiation that diversified from its Cis-Andean sister-group about 4 million years ago (Mya; Abreu et al., 2020a). They are conspicuous inhabitants of the Neotropical rainforests and currently occupy a large geographical area across Amazonia and Atlantic Forest that spans seven of the 13 countries in South America (Abreu et al., 2020b; Vivo & Carmignotto, 2015). The latest taxonomic study investigating species limits within *Guerlinguetus*, based on phenotypic variation, proposed that this genus is composed of two broadly distributed species: *G. aestuans* occurring throughout the Amazonas basin in Brazil, Colombia, and Venezuela, as well as in the Guyana shield (French Guiana, Guyana, and Suriname); and *G. brasiliensis* with a disjunct distribution in the eastern Brazilian Amazonia and in the Brazilian and Argentinean Atlantic Forest (Vivo & Carmignotto, 2015). However, recent molecular studies based on mitochondrial genomes (Abreu et al., 2020b) and Ultraconserved Elements (UCEs; Abreu et al., 2022) both fail to recover reciprocal monophyly for these species and suggested that the species diversity in this genus is likely underestimated. Furthermore, discrepant relationships and distinct species schemes were obtained using mtDNA and UCEs (Abreu et al., 2020b; Abreu et al., 2022), raising questions about the underlaying processes promoting discordant evolutionary histories on the mitochondrial and nuclear genomes and on how this mito-nuclear disagreement affects *Guerlinguetus* species recognition. Therefore, forest-dwelling squirrels of the genus *Guerlinguetus* offer an opportunity to investigate speciation processes and the diversification of an important mammalian radiation across dynamic-changing landscapes, while also contributing to build a broader picture of the biogeographical history of the Neotropical rainforests.

We obtained genomic data (UCEs and single nucleotide polymorphisms) from samples covering most of the geographic range of the genus *Guerlinguetus* and integrated fine-scale population genetic analyses with quantitative molecular species delimitation methods, species tree reconstructions, tests for gene flow, and migration models in a multilayered genomic approach to investigate the diversification of tree squirrels across tropical rainforests of South America. We first discovered population clusters and structure in our sample and then compared the results from different population genetic analyses with the phylogenomic relationships of each specimen to formulate a new hypothesis on species limits. Our new species hypothesis, as well as the species schemes proposed based on phenotypes (Vivo & Carmignotto, 2015) and mitogenomes (Abreu et al., 2020b) were quantitatively compared using species delimitation methods. We further discuss the influence of migration and gene flow when estimating species relationships, and also the putative impacts of these biological phenomena on the observed mito-nuclear discordance. Finally, we explored historical geological and climatic events driving this remarkable tree squirrel speciation while examining new nuances of recent mammalian diversification.

## 2. Materials and Methods

### 2.1 Samples and Data Acquisition

We obtained UCE enriched contigs from 67 specimens of *Guerlinguetus* generated by Abreu et al. (2022), available at GenBank under the BioProject PRJNA847638. Those specimens came from 57 localities, representing a large portion of the known geographic distribution of this genus (Vivo & Carmignotto, 2015; Supplementary Figure S1). In addition, we obtained UCE data from five related taxa (*Tamiasciurus douglasii*, *Sciurus vulgaris*, *Echinosciurus variegatoides*, “*Microsciurus*” *flaviventer*, and *Hadrosciurus spadiceus*) to be used as outgroups in the phylogenetic inferences. Taxonomic identifications follow Abreu et al. (2020b). A complete list of samples analyzed in this study, accompanying museum catalogue and geographic data, is presented in the Supplementary Table S1.

### 2.2 Data Processing and Locus-based DNA Matrices

To generate UCE loci data matrices, we incorporated de novo assembled contigs into the PHYLUCE 1.6 pipeline (Faircloth, 2016; Faircloth et al., 2012), matched those contigs to the uce-5k-probe-set (Faircloth et al., 2012), and aligned and edge-trimmed the identified UCE loci using MAFFT 7 (Katoh & Standley, 2013; Nakamura et al., 2018). We generated a UCE data-matrix by including all loci with at least 50% of sample representativeness using the function “phyluce_align_get_only_loci_with_min_taxa” and subsequently converted this matrix into a phylip format using the function “phyluce_align_format_nexus_files_for_raxml”. Our final UCE data set included 3,642 loci (1,495,138 bp) from 67 individuals of *Guerlinguetus* in addition to five outgroups (Supplementary Data set 1). UCE data processing was done at the Smithsonian Institution High Performance Cluster (SI-HPC; 10.25572/SIHPC).

### 2.3 SNP Calling and Quality Filtering

To assess genetic diversity and structure in *Guerlinguetus*, we extracted single nucleotide polymorphisms (SNPs) from the complete UCE data (4,233 UCE loci, spanning 67 individuals). Different approaches are available to extract SNPs from UCE data (e.g., Andermann et al., 2018; Giarla & Esselstyn, 2015; Parker et al., 2020), each one being advantageous depending upon the level of genomic variation that is intended to be captured. Some methods are preferred when performing across species comparisons while others might be better suited for the analysis of very shallow intraspecific population structure. In this study, we created a unique workflow to phase UCE data following initial steps from the pipeline published by Andermann et al. (2018)—which allowed us to create multiple sequences alignments in a very computationally efficient manner—, and then we phased the reads integrating steps from Giarla & Esselstyn (2015) and Parker et al. (2020). Specifically, we followed the PHYLUCE 1.6 pipeline (Faircloth, 2016) to generate per locus trimmed sequence alignments. These alignments were imported in Geneious R11 (Kearse et al., 2012) where we generated consensus sequences for each locus and subsequently concatenated all sequences to build a pseudogenomic reference, following Parker et al. (2020). To phase UCE data, either individual-specific reference sequences or a pseudogenomic reference is needed to align cleaned raw reads. We tested both strategies and we adopted the pseudogenomic approach because it ultimately allowed us to obtain a larger number of SNPs. We created sequence dictionaries and reference indices from the pseudogenomic reference (which included a total of 1,602,926 base pairs) using SAMtools 1.14 (Danecek et al., 2021). We used BWA-mem algorithm implemented through PHYLUCE (function “phyluce_snp_bwa_multiple_align”) to map cleaned FASTQ read files against the pseudogenomic reference, resulting in indexed BAM files for each sample. SNPs were called with the Genome Analysis Toolkit 4.1.3.0 (GATK; McKenna et al., 2010), following parameters suggested by Giarla & Esselstyn (2015) and Parker et al. (2020). Specifically, we located and excluded indels, called variants, applied quality filters (QD < 2.0; FS > 60.0; MQ < 30.0; HaplotypeScore > 13.0; MappingQualityRankSum < -12.5; ReadPosRankSum < -8.0), and finally selected only biallelic variants, which resulted in a total of 4,570 SNPs. Subsequently, VCFtools 0.1.16 (Danecek et al., 2011) was used to select SNPs with a minor allele count greater than 5 and a minor allele frequency ≥ 5%, generate a set of unlinked SNPs by thinning sites so that no two sites are within 2,000 bp from one another, and to select a database with only 25% of missing data. We also opted to drop three individuals due to their large amounts of missing data, resulting in a SNPs data set with 64 individuals (versus 67 individuals in the UCE loci data set). Additionally, the function HDplot.R (McKinney et al., 2017) was used to identify putative paralogous SNPs by removing those with heterozygosity >0.75 and a D score >10. Therefore, our final SNP data set included 1,169 biallelic, unlinked SNPs from 64 individuals (Supplementary Data set 2). All bioinformatic steps with PHYLUCE and GATK were performed in the SI-HPC (10.25572/SIHPC). Our detailed bioinformatics workflow is described in the Supplementary File 1.

### 2.4 Specimen-based Phylogenetic Analyses

The UCE loci data set (50% matrix completeness; 3,642 loci; Supplementary Data set 1) was analyzed using concatenation and coalescence to infer relationships among all 67 sequenced individuals. For the concatenation approach, we performed ML analyses using RAxML 8.2.7 (Stamatakis, 2014) and IQ-Tree 2.1.1 (Minh et al., 2020). RAxML analysis was performed with the unpartitioned UCE matrix and GTR nucleotide substitution model with gamma-distributed rate heterogeneity (GTRGAMMA). Branch support was accessed from 100 rapid bootstrap replicates and the best-scoring ML tree selected from 10 independent searches. IQ-Tree analysis was performed with the data set partitioned by loci and with GTR + G model of substitution set for each locus. Branch support was assessed from 1,000 ultrafast bootstrap replicates (UFBoot2; Hoang et al., 2018). All RAxML analyses were conducted at the SI-HPC (10.25572/SIHPC) and the IQ-Tree analysis at CIPRES (Miller et al., 2010). For the coalescent approach, we used a site-based method in the software SVDquartets (Chifman & Kubatko, 2014) implemented in PAUP* 4.0a166 (Swofford, 2003). We performed exhaustive quartet sampling—testing all possible quartets—to search for the best-scoring tree. Branch support was obtained from 100 standard bootstrap replicates.

### 2.5 Genetic Structure and Sample Clustering

Genetic structure was evaluated using the data set of 1,169 biallelic, unlinked SNPs (Supplementary Data set 2) with three different approaches: Principal Component Analysis (PCA), Discriminant Analysis of Principal Component (DAPC), and Structure Analysis. PCA and DAPC were used to estimate the number of groups with genetically related individuals. The PCA was performed with the “dudi.pca” function from the R package ade4 (Dray & Dufour, 2007). After the PCA, a Tracy-Widom test (Tracy & Widom, 1994) was performed with the “tw” function from the R package AssocTests (Wang et al., 2020) to select the significant number of PCs that could be interpreted as genetic clusters. In the DAPC the best number of genetic clusters (K) was tested using all PCs (100% of the variance) and Bayesian Information Criterion (BIC) with the “find.clusters” function from the R package adegenet (Jombart & Ahmed, 2011), where the best K is the one with the lowest value, as with the cross-entropy evaluation. The “xvalDapc” function in adegenet package was used to select the best number of PCs to recover genetic clusters (Miller et al., 2020).

The software Structure 2.3.4 (Pritchard & Donnelly, 2000) was used to infer the number of genetic clusters (K), calculate ancestry coefficients for each individual, and assign individuals to ancestral populations. Because the Structure analysis may not necessarily be robust enough to recover a stratified geographic organization (Massatti & Knowles, 2014), hierarchical Structure analysis was also applied to assess substructure within populations. Specifically, analyses were carried out with the entire sample and subsequently for each of the subsets identified as distinct genetic clusters (see Massatti & Knowles, 2014). The data set was analyzed for a K-value from 1-15. Ten independent runs were performed with 500,000 MCMC iterations each and the first 100,000 discarded as burn-in. The software Clumpak (Kopelman et al., 2015) was used to assign individuals to their ancestral history, and the best K was selected using the ΔK method (Evanno et al., 2005) implemented in the software Structure Harvester 0.6.94 (Earl & VonHoldt, 2012).

### 2.6 Quantifying Genetic Differentiation

To examine the genetic differentiation among sample clusters suggested by the Structure analysis, we calculated pairwise weighted *F*_ST_ values (Weir & Cockerham, 1984) using the “gl.fst.pop” function from R package dartR 1.9.6 (Gruber et al., 2018). Moreover, Structure analyses might detect a larger number of genetic clusters in the presence of isolation-by-distance (IBD). To avoid such bias and evaluate whether geographical distance is playing any role on the genetic structure here reported, we conducted Mantel tests (Mantel, 1967) within each genetic cluster represented by samples covering a sizable geographic area (i.e., Northern Amazonia, Eastern Amazonia, and Atlantic Forest). We used the function “mantel.rtest” from the R package ade4 (Dray & Dufour, 2007) and ran 10,000 permutations. The genetic distance matrix was generated with the “mat_gen_dist” function from R package graph4lg (Savary et al., 2021), and the geographic distance matrix was built with the “earth.dist” function from R package fossil (Vavrek, 2011).

### 2.7 Species Tree Estimation and Quantitative Species Delimitation

Based on the consensual results from the analyses described above on the genomic diversity and structure, and phylogenetic relationships, we formulated a new hypothesis on species limits within *Guerlinguetus.* We estimated species trees to infer relationships among the putative species and used quantitative species delimitation methods to examine the statistical support of the proposed species. Species trees were estimated using two coalescent approaches, a site-based (also referred to as a quartet-based) coalescent method in SVDquartets (Chifman & Kubatko, 2014), implemented in PAUP* 4.0a168 (Swofford, 2003), and a locus-based coalescent method in the software Bayesian Phylogenetics and Phylogeography (BPP 4.1.4; Flouri et al., 2018; Yang, 2015), in addition to a hierarchical Bayesian method in the software SNAPP 1.5.2 (Bryant et al., 2012) implemented in BEAST 2.5 (Bouckaert et al., 2019).

In SVDquartets, species trees are inferred considering the most probable quartet resolution at each alignment site (Chifman & Kubatko, 2014). Multispecies sequence alignments might be composed of entire loci or only representative SNPs, which could be either unlinked or correlated (McLean et al., 2022). To run SVDquartets analyses we used both types of data matrices, i.e., the alignment of 3,642 UCE loci (Supplementary Data set 1) and the alignment of 1,169 biallelic, unlinked SNPs (Supplementary Data set 2). These alignments included 67 and 64 specimens of *Guerlinguetus*, respectively, and a tax partition block was inserted to assign those specimens to each of the putative species. The best-scoring species trees were estimated by performing exhaustive quartet sampling, and branch supports were assessed by running 100 (loci) or 1,000 (SNPs) standard bootstrap replicates.

In the second coalescent method, BPP, we analyzed a subset of 500 UCE loci (corresponding to the loci with the highest sample representativeness) and 67 specimens of *Guerlinguetus* (Supplementary Data set 3) assigned to the putative species. We made this subset of loci to reduce computational burden. BPP estimates species trees assuming that gene trees at different loci are independent and accommodating the discordant gene trees (Jiao et al., 2021). BPP species tree (analysis A01) was estimated considering a parameter setting of θ = IG (3, 0.002) and τ = IG (3, 0,004). We ran 100,000 MCMC generations sampling every 10 generations, after a burn-in of 50,000 generations. Finetune parameters adjustments were the same as described above.

Lastly, we used SNAPP to estimate a species tree using SNPs data. To make SNAPP analysis computationally tractable, we had to reduce our original SNPs data set to include up to four samples per putative species, following recommendations in Leaché & Bouckaert (2018). Our SNAPP matrix included 1,155 SNPs from 17 specimens (Supplementary Data set 4). SNAPP uses likelihood algorithm within a Bayesian MCMC sampler; a great advantage of SNAPP over other Bayesian methods is that it bypasses the gene trees and computes species tree likelihoods directly from genetic markers (Bryant et al., 2012). The input file for SNAPP was generated using BEAUTi (Bouckaert et al., 2019) with most default parameters. We only disabled the “Include non-polymorphic sites” option as our SNPs alignment did not include monomorphic sites, and we maintained the log likelihood correction. The forward (u) and reverse (v) mutation rates were set to be calculated by SNAPP. SNAPP analysis was run for 1 million generations, sampling states and trees every 1,000 generations. At the end, we discarded the first 25% of samples as burn-in. Convergence of the runs and adequate effective sample sizes (ESS >200) for all model parameters were verified in Tracer 1.7 (Rambaut et al., 2018). We visualized the sampled trees using DensiTree (Bouckaert, 2010) and summarized them in TreeAnnotator (Bouckaert et al., 2019) using the maximum clade credibility for the target tree.

To evaluate statistical support of the putative species, we first used the Bayesian multispecies coalescent model in the software BPP (Yang, 2015). To run species delimitation in BPP (analysis A10) a guide species tree is required. We used the three distinct topologies that resulted from our species tree analyses (SVDquartets resulted in one topology, BPP in another, and SNAPP in a third topology). Therefore, for each of these three inferred topologies, we performed four species delimitation analyses varying priors of ancestral population size (θ) and root age (τ) [(i) θ = IG (3, 0.2) and τ = IG (3, 0.4); (ii) θ = IG (3, 0.2) and τ = IG (3, 0.004); (iii) θ = IG (3, 0.002) and τ = IG (3, 0.4); and (iv) θ = IG (3, 0.002) and τ = IG (3, 0.004)] and using a subset of 500 UCE loci (Supplementary Data set 3) in BPP 4.1.4 (Flouri et al., 2018). The set of four analyses performed for each distinct topology was repeated three times, allowing us to examine the convergence of MCMC runs. Each BPP analysis was performed by running 300,000 MCMC generations, sampling every 5 generations, after a burn-in of 50,000 generations. We applied an automatic adjustment of the finetune parameters, allowing swapping rates to range between 0.30 and 0.70 (Yang, 2015), and considered Posterior probability (PP) well supported when PP > 0.95.

We also used the Bayes Factor Delimitation with genomic data (BFD*; Leaché et al., 2014) to determine the most plausible species assignment of individual lineages. BFD* method is implemented through SNAPP (Bryant et al., 2012) which runs in the BEAST platform (Bouckaert et al., 2019). For this approach, we do not need to furnish a species tree—as needed for BPP—, but we do need to furnish a priori hypotheses of species limits to be compared. Therefore, we compared three different species-limits hypotheses: (i) two-species, based on the current taxonomic arrangement for the genus *Guerlinguetus* proposed by Vivo & Carmignotto (2015); (ii) four-species, based on molecular species delimitation analyses with mtDNA performed in Abreu et al. (2020b); and (iii) six-species scheme, based on our combined results from population clustering and specimens phylogenetic relationships (same scheme tested with BPP). Our BFD* data matrix included 1,155 SNPs from 17 specimens (same matrix used for the SNAPP species tree; Supplementary Data set 4). For each hypothesis we conducted a path sampling analysis and calculated the marginal likelihood estimates (MLE). For MLE calculation we used 48 steps with 500,000 iterations, a preBurnin of 10,000 iterations, α value of 0.3, and different gamma prior distributions for alpha and beta (2, 20; 2, 2,000; and the default 11.75, 109.73). As the results were similar among different gamma prior distributions for alpha and beta, only the results with the default values are shown. This set of path sampling analysis was repeated three times using different seeds to assess consistency. For all analyses, models were ranked by their MLE, and MLEs were compared using Bayes Factors.

### 2.8 Testing for Gene Flow, Introgression, and Migration Models

To investigate potential sources of conflicts (e.g., gene flow) in our species trees, we used an additional method of species tree estimation that, under a ML framework, progressively infers migration events between pairs of species. We used our data set of 1,169 biallelic, unlinked SNPs (Supplementary Data set 2), including 64 individuals assigned to the six putative species, as input for TreeMix (Pickrell & Pritchard, 2012). TreeMix considers allele frequency covariance to first estimate a species tree with no migration edges and we set the analysis to subsequently add up to five migration edges to the phylogeny. To determine the best-fit model (i.e., no migration or one to five migration events) explaining the variation in our data, we used the R package OptM (Fitak, 2021), which automates the process of selecting the optimal number of migration edges to include. Specifically, we used the Evanno method to calculate an ad hoc statistic (Δ*m*) based on the second-order rate of change in the likelihood weighted by the standard deviation (Evanno et al., 2005). The OptM method requires multiple iterations for each value of M (the number of migration edges), and variation across those interactions for each M must be achieved. Therefore, to obtain different likelihoods for each M, we ran 15 interactions varying the parameter -k (window size in which SNPs will be grouped together to account for linkage disequilibrium) in TreeMix, as per author recommendation (Fitak, 2021). Lastly, we ran 100 bootstrap replicates for the TreeMix species tree with the optimal number of migration edges, and summarized the branch support values on this tree using DendroPy (Sukumaran & Holder, 2010).

Additionally, we tested for hybridization between species lineages of *Guerlinguetus* using the program Dsuite (Malinsky et al., 2021), which also uses allele frequency data from an input of unlinked biallelic SNPs to calculate (i) Patterson’s D (also known as the ABBA-BABA statistic, Durand et al., 2011) and (ii) f4-ratio statistics for all possible species trios with a pre-determined fourth species being used as outgroup. Dsuite also has implemented the f-branch statistic (Malinsky et al., 2018), which can test for excess allele sharing, identifying instances of gene flow between all branches, including internal branches on a specified species tree. We used the SVDquartets species tree, which is identical to the TreeMix tree, as the guide species tree to calculate the f-branch statistic.

Because of the results obtained with TreeMix and Dsuite showing positive signs of migration and an excess of allele sharing between some species lineages, we then used the program Estimated Effective Migration Surfaces (EEMS; Petkova et al., 2016) to spatially visualize the most probable routes of migration but also spatial interruptions (e.g., geographic and ecologic barriers) potentially associated with isolation/divergence among sampled populations. We used as input a matrix of pairwise Euclidean distances constructed using the function “dist.genpop” from R package adegenet (Jombart, 2008). To create this matrix, we first converted our .vcf file with 1,169 biallelic, unlinked SNPs (Supplementary Data set 2) to a genind object using the function “vcfR2genind”, and then transformed it into a genpop object using “genind2genpop” function from adegenet (Jombart, 2008). EEMS also requires as input two files with georeferenced information: the first is a list of geographic coordinates for all sample collecting localities, and the second is a polygon of the geographic area comprising all individual sampling localities. The polygon was manually created and saved as a shape file in QGIS 3.28.5 (https://qgis.org/) to ensure that all sampled localities would be included in the analysis. We selected the deme sizes to 800 and we ran EEMS for 10 million MCMC generations, discarding the initial 25% of the runs as burn-in. We finally verified the convergence of the runs by examining the MCMC trace files.

### 2.9 Divergence times

To estimate historical demographic parameters such as divergence time and migration rates we used G-PhoCS 1.3 (Gronau et al., 2011). G-PhoCS accepts as input a set of multiple sequence alignments and therefore we use the same dataset used in the analyses with BPP. Species were defined according to the six-species hypothesis and historical relationships between them were defined based on the SVDquartets and TreeMix species tree. We performed the analyses with and without migration bands. For the migration analyses, we included migration bands for two species pairs based on the TreeMix results. We estimated for each phylogenetic node θ (theta), τ (tau), and m (migration rates). Theta values were converted to effective population size using θ = 4Neμ. Tau values were converted to divergence time in years with τ = Tμ (where T is the divergence time in generations). The migration rates per generation between species were scaled by the mutation rate with m=MST/μ, where MST is the proportion of individuals in one population that arrived from another population in each generation. For the θ and τ priors we applied gamma distributions (given by parameters shape, α, and rate, β) setting α = 1 and β = 20. For the migration band the gamma distribution was set with α = 0.002 and β = 0.001. We also applied mutation rate of μ = 2.0 × 10^−9^ mutations/site/generation (Gossmann et al., 2019), and to convert estimated coalescent times from the number of generations to years, we assumed a generation time of 1.5 year (Thorington & Ferrell, 2006). Three independent runs were performed for each model using 500,000 MCMC generations, sampling every 100 generations, and discarding the 50,000 initial generations as burn-in. We combined the runs in Tracer 1.7 (Rambaut et al., 2018), where we also evaluated the convergence of the parameters and ESS values.

## 3. Results

### 3.1 Sample Clustering, Specimens Relationships, and Species Discovery

The DAPC analysis suggested that the most likely number of population clusters in our data set is nine, based on the BIC method (Figure 1; Supplementary Figure S2; see also Supplementary Information S2 for detailed description of the geographic ranges of the nine population clusters). The Structure analyses recovered the most likely numbers of unique clusters as K = 2 and K = 6, with ΔK values of 18344.935190 and 9.505451 respectively (Supplementary Table S2; Supplementary Figure S3). Several studies have shown that ΔK method is strongly biased toward K = 2 (e.g., Janes et al., 2017). Therefore, considering such bias, as well as the results from other population genetic analyses and phylogenetic estimations (see below), we consider the second-best K (K = 6) as a more appropriate model to describe the genetic structure in our sample. Nevertheless, we show the proportion of individual membership in each cluster as suggested for both K = 2 and K = 6 (Figure 2). The hierarchical Structure approach detected nine distinct population clusters after three rounds of analyses, where three of the six populations suggested by the previous Structure analysis were split, each into two other populations (Supplementary Figure S4).

**Figure 1.**
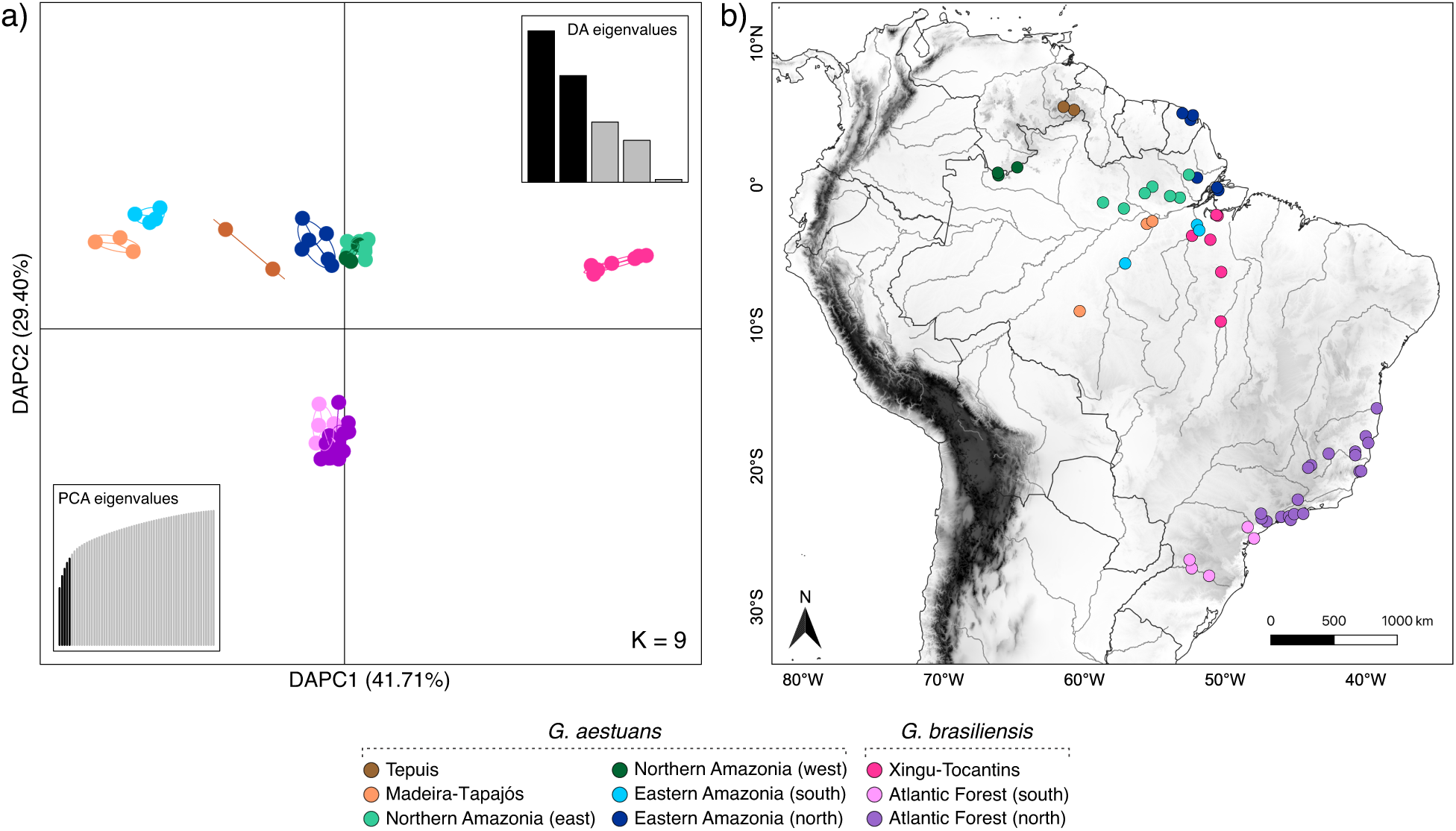
Population clustering as revealed by the Discriminant Analysis of Principal Component (DAPC) using SNP data (1,169 SNPs, 64 individuals). Results are shown for the best scheme (K = 9) as suggested by the Bayesian Information Criterion. **a)** Two-dimensional scatterplot showing samples dispersion in the first two DAPC axes, along with the cumulative eigenvalues from the Discriminant Analysis (top right corner) and the cumulative eigenvalues from the Principal Component Analysis (bottom left corner). **b)** Sampling localities of all specimens included in the DAPC. The colors correspond to the clusters representing K = 9 scheme. The nine population clusters are named according to their geographic location and identified to which species they belong according to the current taxonomy.

**Figure 2.**
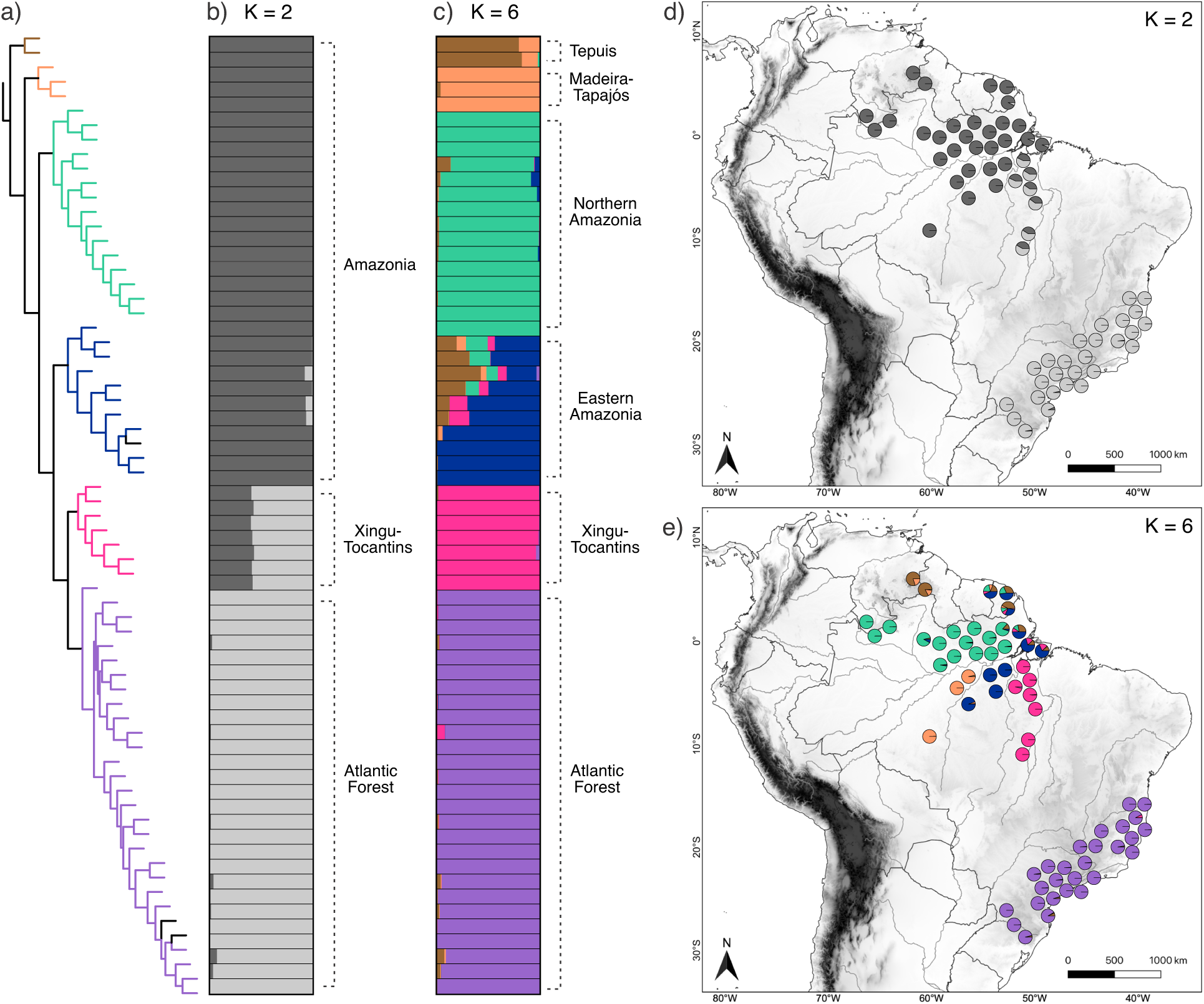
Phylogenetic relationships of *Guerlinguetus* based on UCE data (3,642 loci, 67 specimens) (a), and population structure and ancestry assignments of *Guerlinguetus* based on the Structure analysis of SNP data (1,169 SNPs, 64 specimens) (b, c, d, e). **a)** Phylogenetic relationships as inferred from the coalescent approach in SVDquartets, with correspondence to the bar plots. **b)** Ancestry assignments and genetic admixture shown as bar plots for each individual according to the most optimal clustering scheme (K = 2). **c)** Ancestry assignments and genetic admixture shown as bar plots for each individual according to the second most optimal clustering scheme (K = 6). **d)** Pie charts showing ancestry assignments and genetic admixture for each individual on K = 2 scheme plotted on the map according to the sampling localities. **e)** Pie charts showing ancestry assignments and genetic admixture for each individual on K = 6 scheme plotted on the map according to the sampling localities. Colors on the phylogeny (a) correspond to colors of K = 6 scheme (shown on c and e). The three terminals of the phylogeny colored in black are specimens exclusively analyzed for UCEs but not for SNPs.

Assignments of individuals to unique clusters by the hierarchical Structure and the DAPC were almost identical, with only a few exceptions within the Atlantic Forest sample (see Figure 3). These results differ from the regular Structure analysis, as in the latter, individuals from northern Amazonia, eastern Amazonia, and the Atlantic Forest composed three respectively unique populations, while in the DAPC and in the hierarchical Structure those individuals are further split into six populations, with two subpopulations representing each one of these large geographic regions (Figure 3). The larger number of clusters found by the DAPC and hierarchical Structure might be a response to strong population variation in regions with wide latitudinal (i.e., eastern Amazonia and Atlantic Forest) or longitudinal (i.e., northern Amazonia) gradients. This assumption is partially supported by the Mantel test results, which revealed significant correlation between genetic and geographical distances within the Atlantic Forest cluster (n= 27, r = 0.47, p = 0.000; Supplementary Figure S5) and the Northern Amazonia cluster (n = 15, r = 0.65, p = 0.000; Supplementary Figure S6), but the same result was not found for the second widespread Amazonian cluster, the Eastern Amazonia (n = 10, r = 0.31, p = 0.052; Supplementary Figure S7), suggesting that factors other than geographic distance are responsible for the structuring patterns of the genetic diversity of this group.

**Figure 3.**
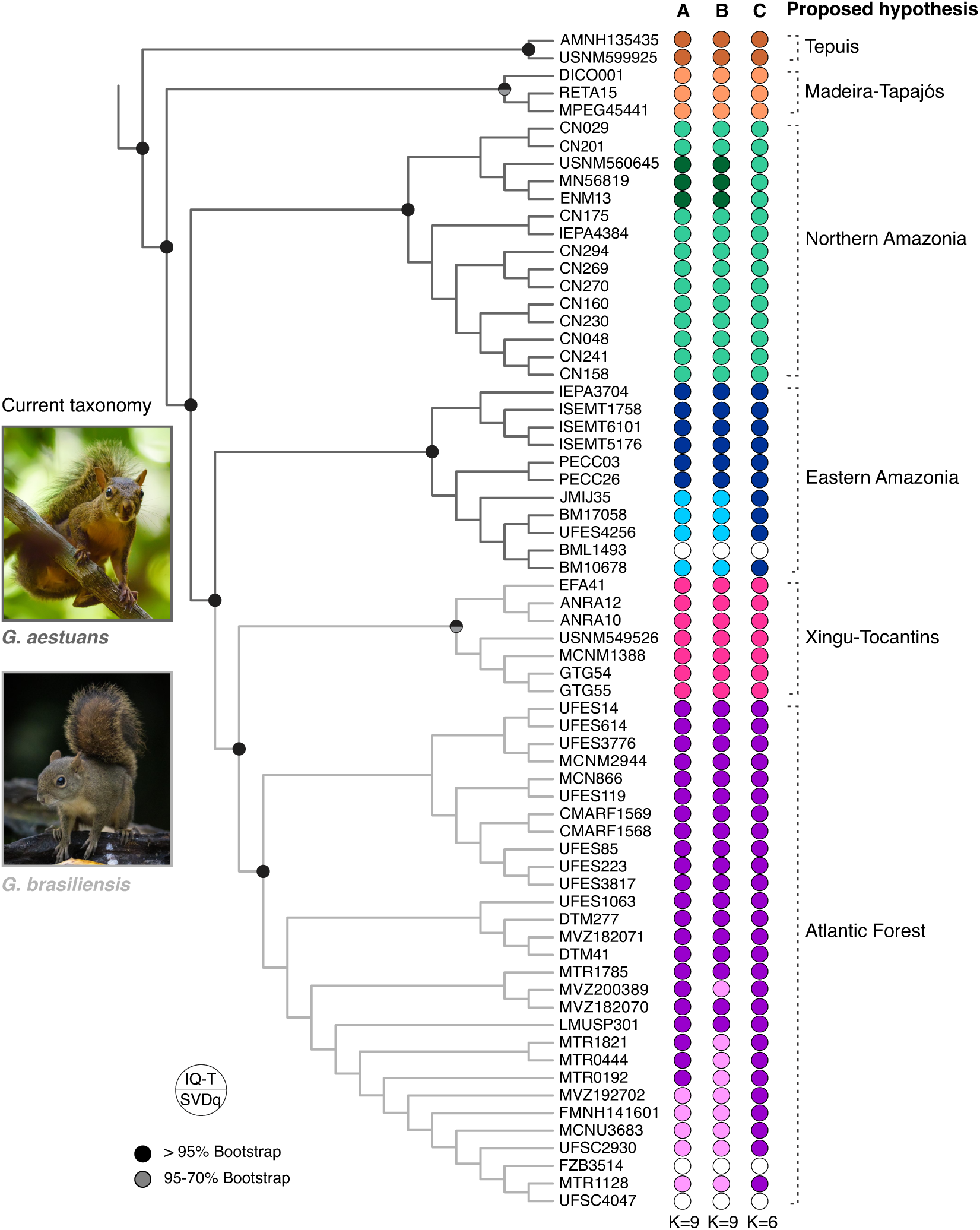
Phylogenetic relationships of *Guerlinguetus* inferred with a maximum likelihood approach in the software IQ-Tree using UCE data (3,642 loci, 67 specimens). The branches on the phylogeny are color coded according to the current taxonomy (Vivo & Carmignotto, 2015): *G. aestuans* (dark gray) and *G. brasiliensis* (light gray). The population clustering using SNPs (1,169 SNPs, 64 specimens) are shown on the right side of the phylogeny, as recovered by the DAPC (A; K = 9), Hierarchical Structure (B; K = 9), and Structure (C; K = 6). The three terminals represented by white circles are specimens exclusively analyzed for UCEs but not for SNPs. Our proposed species hypothesis is identified on the last column, with six lineages named according to geographic range.

Additionally, the DAPC analysis highlighted a strong segregation among populations from the Atlantic Forest, the Xingu-Tocantins interfluve, and the rest of Amazonia (Figure 1). The DAPC1 accounted for around 42% of the variation, and clearly separates the Xingu-Tocantins population from all others, whereas the DAPC2 (30% of the variation explained) segregates the Amazonian populations from those from the Atlantic Forest (Figure 1). This pattern is like the result found for the K = 2 on the regular Structure analysis (Figure 2b, d). Remarkably, in this analysis, individuals from the Xingu-Tocantins area were assigned to the Atlantic Forest cluster, although large mixed ancestry was detected for those. Results from the regular Structure analysis (K = 6; Figure 2c, e) reaffirmed the segregation of populations from the Atlantic Forest and from the Xingu-Tocantins interfluve (hereafter Atlantic Forest and Xingu-Tocantins groups). This analysis also suggested that the remaining Amazonian populations are split into four distinct groups: one group including two individuals from the highlands of the Guiana Shield (hereafter Tepuis), a group composed of three individuals from southwestern Amazonia (hereafter Madeira-Tapajós), a group of 15 individuals spanning a large area on the northern bank of the Amazonas River, including both west and east of Negro River (hereafter Northern Amazonia), and the last cluster composed of 10 individuals from eastern Amazonia, including localities from both north and south banks of Amazonas River (hereafter Eastern Amazonia). Individuals from the Eastern Amazonia cluster are the ones with the largest amounts of admixture (Figure 2c, e).

Our phylogenetic reconstructions based on loci (data set including 3,642 UCEs and 67 individuals of *Guerlinguetus*) produced topologically similar results using both concatenation (RAxML and IQ-Tree) and coalescence (SVDquartets) (Supplementary Figures S8-10). All methods estimated well-supported, reciprocally monophyletic groups corresponding to the K = 6 Structure scheme (Figure 3). Neither K = 2 (from the regular Structure) nor K = 9 (from DAPC and hierarchical Structure) schemes correspond to reciprocally monophyletic groups on the estimated phylogenies (Figure 3). Only the subdivision of the Northern Amazonia cluster into two groups, as suggested by the DAPC and the hierarchical Structure (see Figure 1 and Supplementary Figure S4), was recovered as reciprocally monophyletic in the SVDquartets topology (Supplementary Figure S10). Therefore, integrating the population structure assignments with the evolutionary relationships of individuals (summarized in the Figure 3), we hypothesized that six putative species might be masked under the current taxonomy of only two species within the widespread genus *Guerlinguetus*, and this new hypothesis was tested in the downstream analyses. We also assessed the genetic differentiation between these six putative species using the *F*_ST_ metric and we observed high pairwise fixation indices (> 0.38) in all comparisons (Table 1). This supports a scenario of deep genetic segregation among them and reinforces our new hypothesis on the species diversity of *Guerlinguetus*.

**Table 1.** Pairwise weighted *F*_st_ values between each of the six putative species of *Guerlinguetus*, delimited in this study.

|  | Tepuis | Madeira-Tapajós | Northern Amazonia | Eastern Amazonia | Xingu-Tocantins |
| --- | --- | --- | --- | --- | --- |
| Madeira-Tapajós | 0.70 |  |  |  |  |
| Northern Amazonia | 0.40 | 0.42 |  |  |  |
| Eastern Amazonia | 0.41 | 0.45 | 0.38 |  |  |
| Xingu-Tocantins | 0.66 | 0.68 | 0.53 | 0.49 |  |
| Atlantic Forest | 0.72 | 0.73 | 0.67 | 0.65 | 0.48 |

### 3.2 Species Relationships and Quantitative Support

Prior to evaluating the quantitative support for the recognition of six distinct species within *Guerlinguetus*, we used different species-tree approaches to estimate the relationships among the hypothesized species. These analyses needed to be done before running the multispecies coalescent models because guide species trees are recommended for more robust species delimitations.

We obtained three distinct topologies using SVDquartets, BPP, and SNAPP. With SVDquartets, the two analyses (one using the data set of 3,642 UCE loci and another using the SNPs data set) retrieved the same topology (Figure 4a). The BPP inference, using the SNPs data set as input, provided a topology very similar to the SVDquartets topology, except for the sister relationship between the Northern and Eastern Amazonian clades that was not recovered with SVDquartets (Figure 4b). The SNAPP analysis provided the most distinctive topology, with an initial split separating the Atlantic Forest + Xingu-Tocantins clade from a clade including all remaining Amazonian species (Figure 4c). Species relationships within the Amazonian clade also differ from those inferred with the other two methods. The simultaneously visualization of all trees sampled by SNAPP reaffirms uncertainties surrounding the phylogenetic positions of the Northern and the Eastern Amazonian groups.

**Figure 4.**
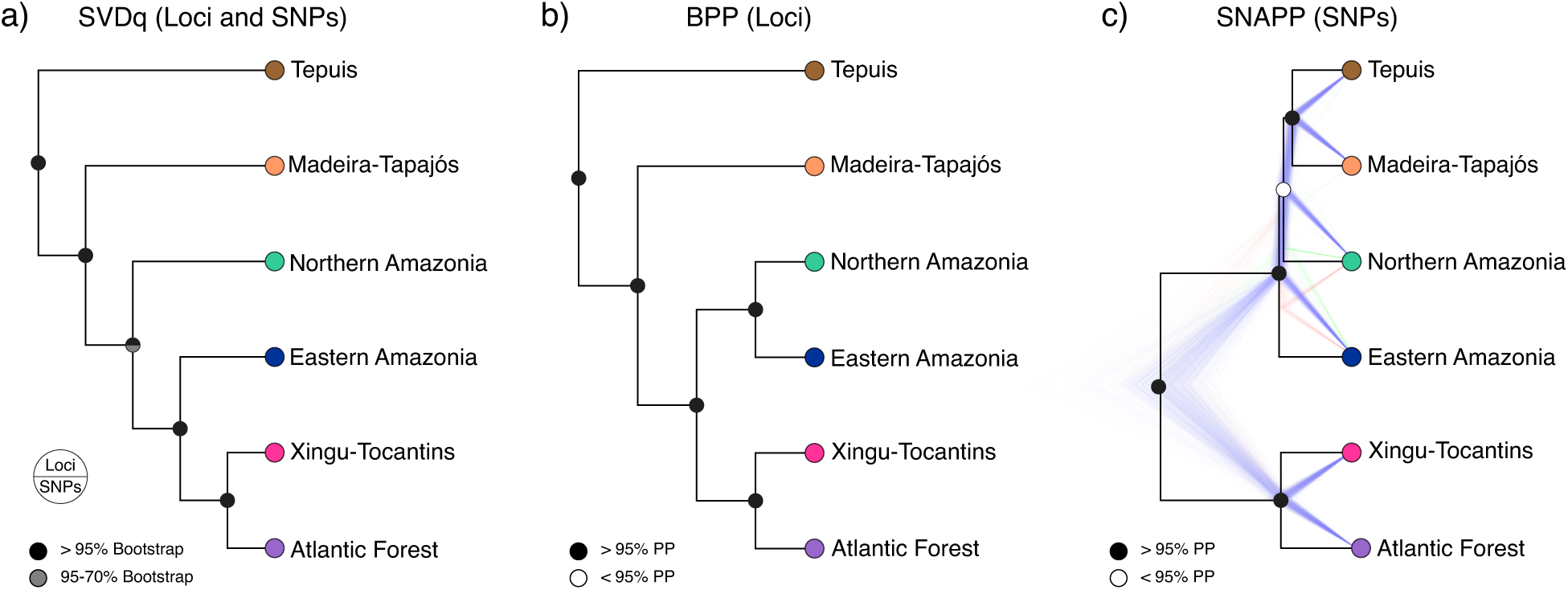
Species trees inferred by assigning all *Guerlinguetus* individuals into six putative species according to our new hypothesis. **a)** Species tree inferred with the site-based coalescent approach in SVDquartets using two different data sets: a data set of 3,642 UCE loci and a data set of 1,169 SNPs. **b)** Species tree inferred with the multispecies coalescent method in BPP using a reduced UCE dataset with 500 loci. **c)** Species tree (maximum clade credibility) inferred with the hierarchical Bayesian method in the software SNAPP plotted over 1,000 trees sampled via MCMC from the posterior distribution of species trees.

These three different topologies were subsequently tested with multispecies coalescent models in BPP, and full support (PP = 1) was obtained for the recognition of the six putative species in all distinct scenarios of ancestral population size and root age assumed. The quantitative molecular species limits using the BFD*, which tested three distinct species schemes (two-species, four-species, and six-species), indicated the six-species scenario as the best scoring according to the Bayes Factors ranking (Table 2).

**Table 2.** Results of BFD* analyses testing the support of competing species limits hypotheses for *Guerlinguetus*. For each species hypothesis scheme, number of species, marginal likelihood estimates (MLE), Bayes factor (calculated as BF = 2*(model1 - model2) and rank are shown. As different gamma prior distributions for alpha and beta retrieved similar results, only the results using default parameter are shown.

| Species hypothesis | N Species | MLE | Bayes factor | Rank |
| --- | --- | --- | --- | --- |
| Morphology* | 2 | -12617.571 | 0 | 3 |
| mtDNA** | 4 | -11633.439 | -1968.26 | 2 |
| UCEs and SNPs (This study) | 6 | -10515.056 | -4205.03 | 1 |

\*According to Vivo & Carmignotto (2015)
\*\*According to Abreu et al. (2020b)

### 3.3 Gene flow and Migration

Although we obtained strong support to accept our hypothesis of six species within *Guerlinguetus*, we were not able to unequivocally estimate their relationships with the approaches presented above. One reason for that could relate to the fact that those methods assume no (or negligible) gene flow. Therefore, to investigate potential violations of this assumption, we estimated another species tree using a framework that progressively adds migration events to explain the overall variance in allele frequencies. Based on the Δ*m* values obtained for six models tested with TreeMix, the model including two migration edges better explained the variation in our data (Figure 5a; Supplementary Figure S11). The first estimated migration edge connects the Madeira-Tapajós populations to the nearby Eastern Amazonian populations, and the second connects the Tepuis population to the geographically distant Atlantic Forest populations. However, both migration edges, especially the second, exhibited mid to low weights.

**Figure 5.**
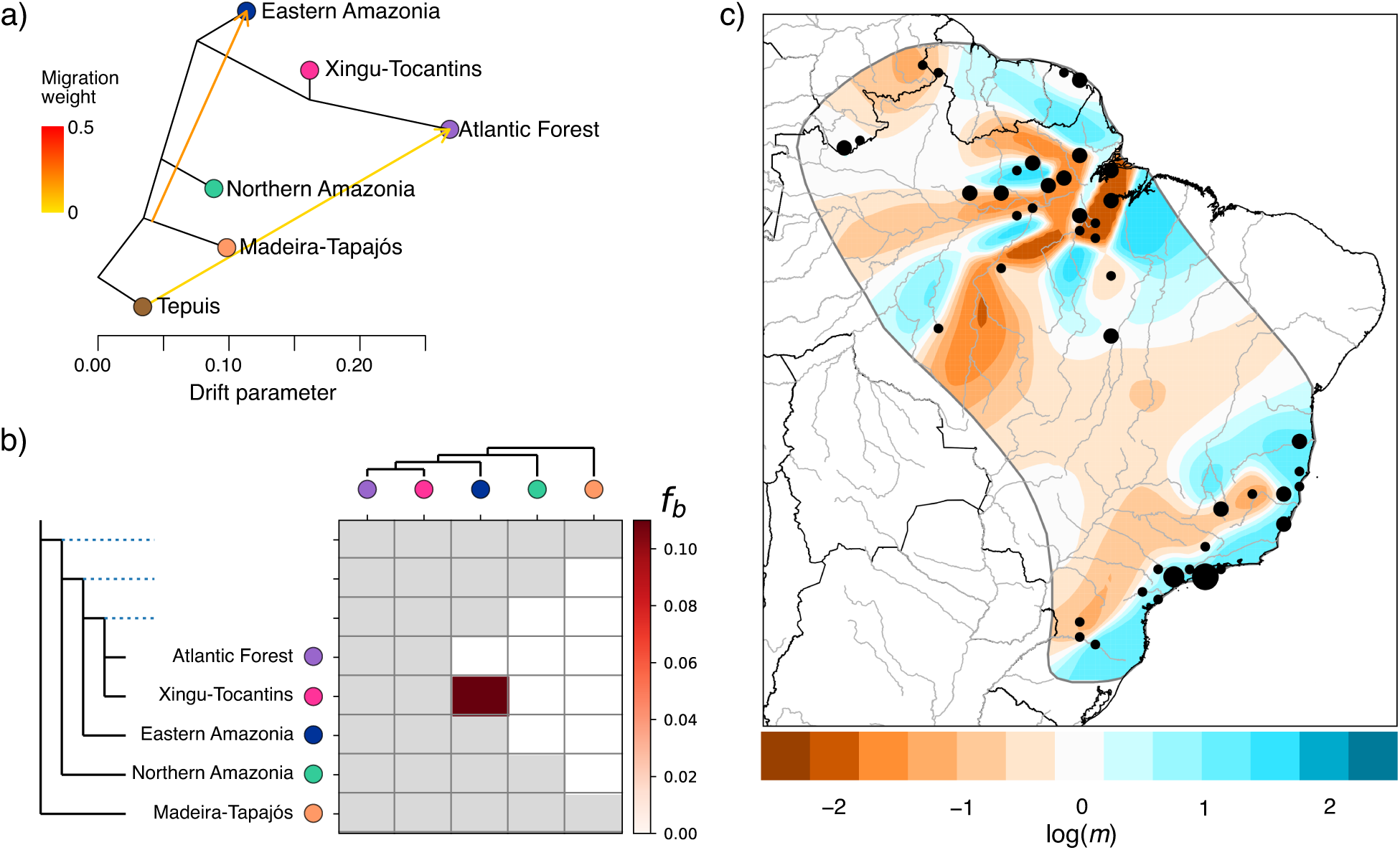
Different approaches for testing for gene flow and migration in the *Guerlinguetus* samples using the SPN data set. **a)** Maximum likelihood tree inferred with TreeMix showing two migration edges as suggested by the Evanno method as the optimal number of migration events explaining the variance in allele frequencies. Branch support was accessed from 100 bootstrap replicates and all branches, except for the Northern Amazonia (bootstrap = 93%), presented statistical support above 95%. **b)** Heatmap showing the values of the f-branch statistic estimated with Dsuite using the topology recovered by TreeMix and SVDquartets. Gray boxes indicate that given branches cannot be tested under the [(A,B),C,D] framework according to the guide topology. The red scale indicates from no allele sharing (white) to excess in allele sharing (dark red). **c)** Geographic representation of effective migration surfaces for the entire *Guerlinguetus* sample, highlighting deviations from a null model of no migration. Effective migration is shown using the logarithm with base 10 scale, where a value of –1 represents a 10–fold decrease in effective migration rate.

Complementarily, we conducted statistical tests for gene flow among trios of species based on the Patterson’s D and f4-ratio statistics using Dsuite and providing the species tree estimated with SVDquartets and TreeMix as the guide tree. We found a single significant (D = 0.172; p < 0.05; Supplementary Table S3) introgression event between the Eastern Amazonia species and the Xingu-Tocantins species. This result was corroborated by an excess in allele sharing detected by the f-branch statistic between those species (Figure 5b).

However, estimating effective migration surface among all populations of *Guerlinguetus*, we observed that the Xingu-Tocantins populations are in about two orders of magnitude deviating from a continuous, positive migration rate with the nearby populations from the Eastern Amazonia species (Figure 5c). Moreover, between the geographic range of the Eastern Amazonian populations and the Madeira-Tapajós populations on the south bank of the Amazonas River−for which TreeMix inferred a migration event−, we detected very low effective migration surfaces, deviating about two orders of magnitude from the average. Remarkably, between the Eastern Amazonian populations from the southern and northern banks of the lower Amazonas River, EEMS showed positive migration rates, supporting all other results indicating that these population represent a single evolutionary unit. Additionally, the EEMS analysis highlighted a strong migration barrier along the Amazonas River (except closer to the river mouth), and it also appears that the migration breaks detected on the south bank of this river coincides at least partially with major riverine tributaries, e.g., Xingu and Tapajós rivers. On the north bank of the Amazonas River, EEMS estimated low migration surfaces in the geographic area limiting the Eastern Amazonia and the Northern Amazonia groups, near the Jari River basin. Likewise, a large area of reduced migration was estimated in the highlands of the western portion of the Guiana Shield. Lastly, a mostly continuous migration surface was detected in the Atlantic Forest, with only lower effective migration rates along the interior, less mesic portions of the Atlantic Forest.

### 3.4 Timing of Diversification

Divergence times estimated without migration bands suggested that the Tepuis species diverged from the other lineages during the mid-Pleistocene (around 1.87 Mya), followed by the divergence of the Madeira-Tapajós lineage around 1.15 Mya. All other diversification events took place in the past 500,000 years. The most recent diversification, between the Xingu-Tocantins and Atlantic Forest lineages, happened about 425,000 years ago (Figure 6; Supplementary Table S4). We could not measure the magnitude of the migration between the Tepuis and Atlantic Forest lineages nor between the Madeira-Tapajós and Eastern Amazonia because migration parameters did not converge even after testing a variety of gamma distributions for the migration priors. This may be due to the negligible amount of migration between these lineages, although it was detected in the TreeMix analysis.

**Figure 6.**
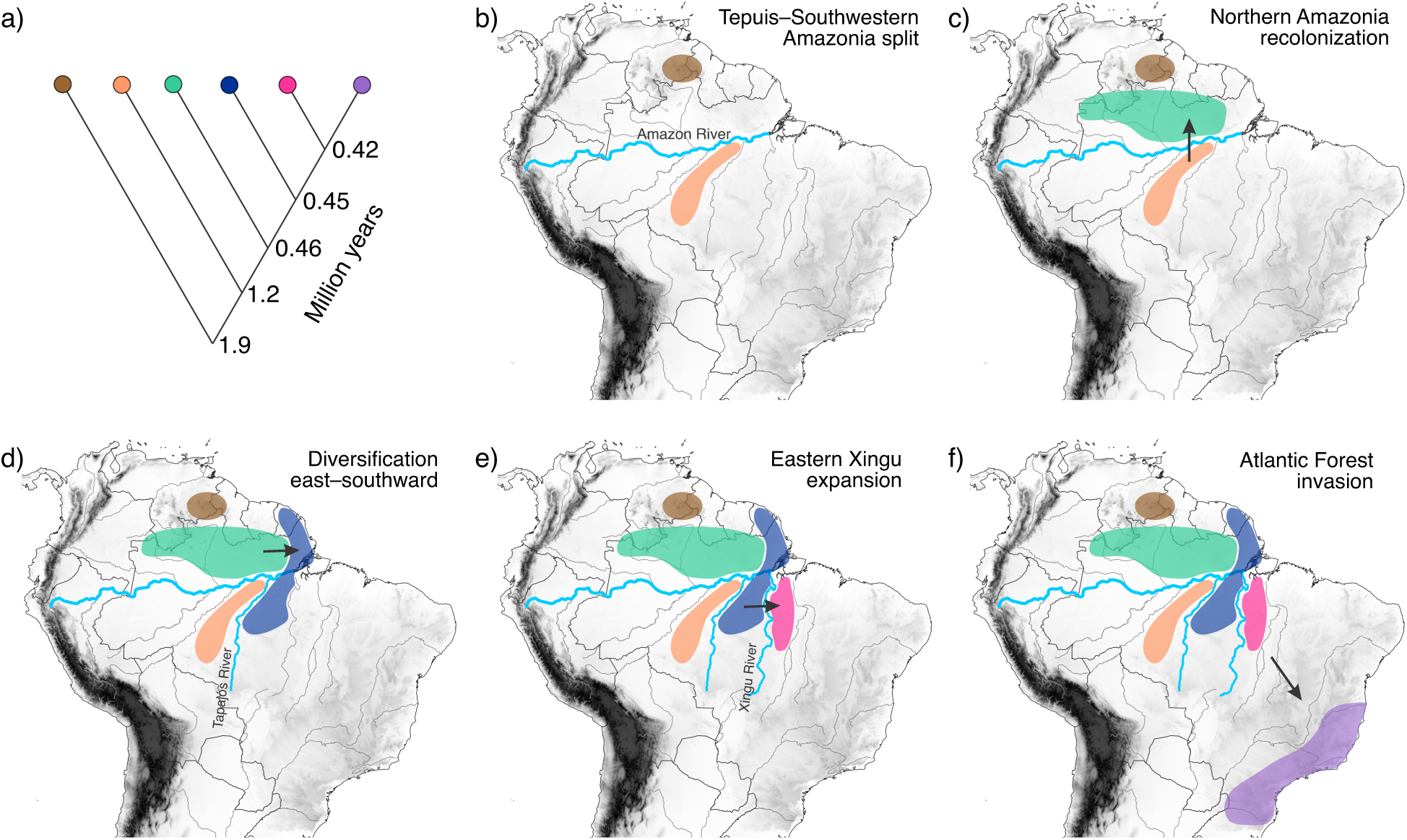
Consolidated diversification hypothesis for tree squirrels of the genus *Guerlinguetus* across tropical rainforests of South America in the past 2 million years. **a)** Diversification times for the six putative species estimated with the demographic G-PhoCS approach. **b)** First split within the genus separating the Tepuis lineage from the Madeira-Tapajos linage on opposite sides of the Amazonas River. **c)** Recolonization of the northern bank of the Amazonas River by the Northern Amazonia lineage. **d)** Expansion to northeastern Amazonia and colonization of the Tapajos-Xingu interfluve (in the south bank of the Amazonas River) by the Eastern Amazonia lineage. **e)** Diversification of the Xingu-Tocantins lineage. **f)** Colonization of the Atlantic Forest on the east coast of South America.

## 4. Discussion

Our results show that phasing and calling SNPs from thousands of UCE loci and incorporating these data in a multilayered approach that combines fine-scale population genetic analyses, quantitative species delimitation, species tree estimation, tests for gene flow, migration and demographic models, as well as time estimation can help to unravel evolutionary trajectories, clarifying complex speciation processes to accurately delimit species and understand biogeographic histories. Specifically, using this comparative approach, we uncovered unexpected species diversity of keystone mammals that are critical in tree-seed predation and dispersion in one of the most fragile and threated ecosystems of the world, the tropical rainforests of South America. We demonstrated that the current taxonomy of tree squirrels of the genus *Guerlinguetus* (recognizing only two broadly distributed species: *G. aestuans* and *G. brasiliensis*; Vivo & Carmignotto, 2015) greatly underestimates the species diversity in this group. The main findings of our study were: (i) population genetic and species delimitation analyses agreed that the most plausible number of species within *Guerlinguetus* is six, contrasting with both the classic morphological revision published by Vivo & Carmignotto (2015) and the mitochondrial phylogeny published by Abreu et al. (2020b); (ii) although we consensually estimated the species limits within *Guerlinguetus*, incongruencies on species relationships still persist, which might be a response to population migration and gene flow taking place in lowlands of eastern Amazonia; and (iii) effective migration surfaces detected important geographic barriers associated with the major Amazonian riverine systems and the mountain ranges of the Guiana shield. In the following sections we address these major findings in detail.

### 4.1 Remarkable species diversity and the power of a multilayered genomic approach to clarify species limits in complex systems

Our results indicate that the genus *Guerlinguetus* harbors at least six distinct species, instead of two (Vivo & Carmignotto, 2015) or four (Abreu et al., 2020b) as previously suggested. This is remarkable but not entirely surprising considering the persistent high rates of mammal species discovery (both new taxa and taxonomic splits) in tropical regions of the globe (Ceballos & Ehrlich, 2009; Parsons et al., 2022). Our results also highlight the value of a comparative approach wherein population genetic analyses and species delimitation methods provide a solid base to clarify taxonomic and evolutionary uncertainties that had been left unresolved by conventional approaches.

In the last taxonomic revision of this group, Vivo and Carmignotto (2015), using morphological information, recognized all populations from the Amazonia (excepting those from east of the Xingu River) as comprising a single species, *G. aestuans*. However, they did acknowledge that there was some degree of morphological uniqueness associated with the Madeira-Tapajós populations and, thus, they allocated those individuals in a subspecies (*G. aestuans gilvigularis*) separated from the remaining *G. aestuans* populations (included in the subspecies *G. aestuans aestuans*). Interestingly, the Madeira-Tapajós populations were, until recently, considered as a distinct species (see Thorington et al., 2012). Our results seem to validate the status of species for the Madeira-Tapajós populations, which were recovered as highly distinct (*F*_st_ ranging from 0.42 to 0.73; Table 1) and with no signs of gene flow (Supplementary Table S3; Figure 5b). Our results also suggest that *G. aestuans aestuans* sensu Vivo and Carmignotto (2015) might be further split into three species, represented by the Tepuis, Northern Amazonia, and Eastern Amazonia clusters. Vivo & Carmignotto (2015) highlighted that samples from highlands of Guiana and Venezuela (correspondent to the Tepuis cluster in our analyses) exhibit morphologic differences from the lowland Amazonian populations, including the presence of postauricular patches. There are several names available in the literature that could potentially be applied to these groups, and a thorough taxonomic investigation is necessary to correctly recommend their application.

Vivo and Carmignotto (2015) also recognized their second species of *Guerlinguetus* to be polytypic, including three subspecies: *G. brasiliensis brasiliensis*, *G. b. ingrami*, and *G. b. paraensis*. Among them, *G. b. ingrami* is represented in our analyses by the Atlantic Forest cluster and all of our results point to the recognition of this group as a full species. The Atlantic Forest cluster is one of the most distinctive in the DAPC analysis (Figure 1) with almost no admixture with the Amazonian groups (Figure 2), which reflects in high *F*_st_ values (Table 1). A similar resolution might be appropriated to *G. b. paraensis*, represented in our analyses by the Xingu-Tocantins cluster (see Figures 1 and 2; Table 1), although Vivo & Carmignotto (2015) had included in their concept of *paraensis* samples from the left bank of Xingu River that we assign to the Eastern Amazonia lineage. Unfortunately, we were not able to include representatives of *G. brasiliensis brasiliensis* (distributed in northeastern Brazil, northwards from the States of Bahia to Ceará; Vivo and Carmignotto, 2015), which prevented us from testing its genetic affinities. An interesting aspect of the species delimitation performed by Vivo and Carmignotto (2015) was the inclusion of populations from the Xingu Tocantins interfluve in their concept of *brasiliensis* (a predominantly Atlantic Forest group) rather than to *aestuans* (an Amazonian radiation). Some of our analyses also suggest a proximity between Atlantic Forest and Xingu Tocantins groups, as the Structure analysis that recovered a K = 2, and the species trees that recovered a sister relationship between these groups.

Two main findings arose by comparing our results with the mtDNA-based molecular species delimitation published by Abreu et al. (2020). First, three mitochondrial groups correspond to putative species as revealed by our multilayered genomic approach, e.g., Tepuis, Madeira-Tapajós, and Atlantic Forest. However, the remaining three putative species recovered here do not represent reciprocally monophyletic groups in the mtDNA phylogeny, e.g., Northern Amazonia, Eastern Amazonia, and Xingu-Tocantins. Specifically, individuals from the Northern and Eastern Amazonian groups were found nested together in the mitogenomic analyses, and specimens composing the Xingu-Tocantins cluster were split in two non-sister clades (Abreu-Jr et al., 2020b). This discordance between the nuclear and mitochondrial evolutionary relationships is addressed below.

In summary, a main take away from these empirical results is that integrating multiple analytical genomic methods can help clarify controversial species limits proposed with base on a single line of evidence, for instance phenotypic data−which might be largely subject to pervasive convergence− or mtDNA sequences−which do not include information from independent loci, represent a uniparental inheritance, and can be rapidly affected by selection pressures (Edwards et al., 2005). Moreover, as acknowledged by DeRaad et al. (2022), integrating a diversity of genomic analytical approaches to delimit species is advantageous to avoid biased assumptions or statistical nuances of any individual approach that might greatly impact the resulting delimitation.

Being able to elucidate species limits and accurately estimate species diversity is especially critical in ecosystems undergoing rapid and permanent changes. Global tropical forests, which are home to at least two-thirds of the world’s biodiversity, are losing species at unprecedent rates (Giam, 2017; Morris, 2010). Two of the six putative species recognized in our study, Xingu-Tocantins and Eastern Amazonia, are distributed in the so called “arch of deforestation” along the southeastern portion of Amazonia (Diniz et al., 2013), amplifying the urgency of describing the diversity of these keystone mammals, particularly involved in the dispersal and maintenance of several native palm trees (Mendes et al., 2019).

### 4.2 Migration and gene flow challenge species tree estimation and likely underlie mito-nuclear discordances

Despite confidently estimating the species-level diversity in *Guerlinguetus*, we struggled to unequivocally infer the relationships among the putative species, as distinct inference methods provided distinct resolutions, particularly regarding the phylogenetic affinities of the Northern and Eastern Amazonian species (Figure 4). Phylogenetic uncertainties might derive from methodological artifacts, such as incomplete taxonomic sampling, inappropriate choice of markers, inadequate selection of models and/or optimality criteria (Giarla & Esselstyn, 2015; Huang et al., 2010). We attempted to overcome analytical biases by estimating species trees using different data sets (loci alignments and SNPs matrices) and employing distinct inference methods (maximum likelihood [TreeMix], coalescence [SVDquartets and BPP], Bayesian [SNAPP]). Using maximum likelihood in TreeMix and a site-based summary coalescent approach in SVDquartets, we recovered the same branching history. However, the multispecies coalescent model in BPP uniquely estimated a sister relationship between the Northern and Eastern Amazonian species, and the Bayesian method in SNAPP uniquely recovered an initial split separating Tepuis+Madeira-Tapajós+Northern Amazonia+Eastern Amazonia from Xingu-Tocantins+Atlantic Forest. Although these discrepant results were nonrecurrent (i.e., each alternative scenario was inferred only once), this might advocate for methodological caveats on phylogenetic estimations with BPP and SNAPP. This is precisely because we had to use reduced input data sets to make these analyses computationally tractable (see Methods section). Also, the fact that these uncertainties mostly involved the Northern and the Eastern Amazonian clades raises additional questions on their nature. Specifically, specimens from these two clades were also found nested together in the mitogenomic hypothesis previously published for the genus (Abreu et al., 2020b). Therefore, uncertainties on the branching history coupled with a mito-nuclear discordance might provide stronger support for additional biological phenomena impacting the evolutionary history of this group rather than methodological artifacts.

Difficulties in estimating species relationships, as well as recurrent mito-nuclear discordances are usually associated with introgression, ILS, and the tempo (i.e., ancient versus recent) and duration (i.e., rapid versus slow) of diversification events (Cai et al., 2021). Mito-nuclear discordance in phylogenetic estimation has become widely observed in several clades of the Tree of Life (Andersen et al., 2021; Good et al., 2008; Kearns et al., 2014; Linnen & Farrell, 2007; Platt et al., 2018), and is currently challenging researchers to understand its nature and its impacts on the delimitation of evolutionary units.

Our results did not show widespread signs of nuclear introgression among the six putative species, but they did show significant introgression and excess in allele sharing between the Eastern Amazonia and the Xingu-Tocantins clusters (Figure 5b; Supplementary Table S3).

Our results also suggested a migration event with stronger weight from the Madeira-Tapajós to the Eastern Amazonia lineage. Therefore, signatures of gene flow and migration particularly along the eastern Amazonia could trace back to past or on-going hybridization events. Even if not extensively detected in the nuclear genome, hybridization might result in larger impacts within the mitochondrial genome, for instance through single or multiple events of mitochondrial capture. In fact, individuals of a given species exhibiting mtDNA from another but having no or little nuclear introgression is a common pattern observed in natural systems (Andersen et al., 2021; Pons et al., 2014). This condition of mitochondrial capture can be very extreme, resulting even in genome replacement (Bonnet et al., 2017), and could be a response to adaptive processes, neutral or nonadaptive captures (including range expansion and hybridization by secondary contact), occasional gene flow, and cryptic speciation (Andersen et al., 2021; Bonnet et al., 2017; O’Connell et al., 2021).

The *Guerlinguetus* mode of speciation with two lineages diverging in allopatry much earlier (in the Plio-Pleistocene border) and the remaining four lineages diversifying very quickly and more recently (mid-late Pleistocene; see next topic for timing and mode of diversification) in subjacent areas throughout mostly the eastern Amazonia, provided opportunity for cryptic speciation and ventures for secondary contact and gene flow through range expansions. Mitochondrial capture events could therefore explain both the mito-nuclear discordance and the recalcitrant nuclear relationships among *Guerlinguetus* species.

Additionally, one could argue that the mito-nuclear discordance found in tree squirrels might just demonstrate weaker phylogenetic signal in the mitochondrial genome in comparison to nuclear markers. This would be expected as differences in the inheritance and recombination patterns of the two genomes can provide distinct levels of phylogenetic signals in their sequences. It is well known that the nuclear genome usually has stronger phylogenetic signal than the mitochondrial genome, because it is affected by both vertical (ancestor-descendant) and horizontal (recombination) gene transfer (Degnan & Rosenberg, 2009; Galtier et al., 2009; Philippe et al., 2011). Moreover, the mitochondrial genome evolves at a faster rate than the nuclear genome due to its smaller size, higher mutation rate, and lack of recombination (Brown et al., 1979; Nabholz et al., 2009). This faster evolution can cause the mitochondrial DNA sequences to become saturated with mutations, which could have also helped to blur the phylogenetic signal in the mitogenomes of *Guerlinguetus* tree squirrels.

### 4.3 Recent speciation in tree squirrels reinforces the young evolutionary history of Amazonian biota

Tropical rainforests of South America, specifically in Amazonia, are home to some of the most diverse biotas on earth (Mannion et al., 2014). Yet, scientists still struggle to answer central ecological and evolutionary questions on the origin and mechanistic drivers of the impressive Amazonian species diversity. Decades of cumulative research have suggested a set of processes governing the origins and diversification of the autochthonous fauna, including biotic diffusion (i.e., geodispersal) across landscapes, dispersal across pre-existing barriers, and isolation and differentiation through vicariance (Cracraft et al., 2020). Among them, vicariance resulting from the riverine system formation or climatic-driven habitat fragmentation appears to be the most recurrent model of diversification for Amazonian vertebrates (Alfaro et al., 2015; Boubli et al., 2015; Pirani et al., 2019; Ribas et al., 2012; Silva et al., 2019). However, a considerable portion of the evolutionary and biogeographic studies of Amazonian vertebrates derives from avian research and, among mammals, this area of research is essentially biased towards primates. Understanding the spatial and temporal evolutionary history of Amazonian biota is, therefore, compromised by the lack of studies spanning other branches of the Tree of Life. Cracraft et al. (2020) advocated that individual, taxon-focused approaches (e.g., phylogeographic analysis of small groups) are important, and authors should contextualize the results within the broader perspective of how organisms evolved across the dynamic Amazonian landscape.

Our results for a clade of widespread tree squirrels, consistently suggest that the first lineage to diverge within this group, around 2 Mya, inhabit highland forests of the Guiana Shield. The second lineage to diverge, about 1.2 Mya, occurs in the southern bank of the Amazonas River, between the Madeira and Tapajós rivers, coincident with the Rondônia area of endemism of Cracraft (1985). The subsequent diversification events took place in a very short temporal window, from 0.46 to 0.42 Mya, and led to (i) the colonization of the lowland forests of the northern bank of the Amazonas River (Northern Amazonia clade); (ii) the eastern Amazonia in both north bank and south bank of the Amazonas River, although in the southern bank this lineage is restricted to the Tapajós-Xingu interfluve, coincident with the Tapajós area of endemism (Ribas et al., 2012); the southeastern Amazonia between the Xingu and Tocantins rivers (Xingu-Tocantins clade), coincident with the Xingu center of endemism (Ribas et al., 2012); and lastly the Atlantic Forest (Atlantic Forest clade), coincident with the Serra do Mar center of endemism (Cracraft, 1985) (Figure 6).

The timing and sequence of these events, i.e., earliest split occurring between the Guiana Shield and the westernmost populations from the Brazilian Shield, followed by colonization events toward southeastern Amazonia, coincide with the diversification history of many upland (“terra firme”) bird lineages (Carneiro et al., 2022; Silva et al., 2019). This pattern also largely mirrors the diversification of spiny rats (genus *Proechimys*; Dalapicolla et al., 2023) and squirrel monkeys (genus *Saimiri*; Alfaro et al., 2015). Specifically, tree squirrels and squirrel monkeys seem to have followed similar pathways for the colonization of eastern Amazonia along the southern bank of the Amazonas River; and for both taxa a dispersal event leading to the recolonization of the northern bank of the Amazonas River appears to have occurred after the initial colonization of southwestern Amazonia (Alfaro et al., 2015).

Reconstructions of the Amazonian vegetation dynamics over the past 1.8 Mya largely agree with the Neotropical tree squirrel diversification history. Between 1.8-1.0 Mya, rainforest vegetation, along with extensive riparian wetland vegetation, dominated the Amazonian landscape, especially in western Amazonia where there was a larger concentration of wetland ecosystems (Hoorn et al., 2010); after 1.0 Mya, contractions of both forests and wetlands started taking place; and from around 400 Kya, about the timing of most speciation events within *Guerlinguetus*, stronger synchrony between changes in vegetation and glacial-interglacial climate cycles is reported (Kern et al., 2023). During this last period, in warmer interglacial times the vegetation was dominated by lowland tropical rainforests, and in cooler glacial periods tropical seasonal forests expanded towards the eastern Amazonia (Kern et al., 2023), coinciding with the *Guerlinguetus* colonization of eastern South America (Figure 6).

The diversification of *Guerlinguetus* tree squirrels also supports the well-established hypothesis of forested connections between the Amazon and the Atlantic forests (Costa, 2003; Costa & Leite, 2012). Our results indicated that the Atlantic Forest species diverged from the Xingu-Tocantis region about 0.4 Mya (Figure 6). Amazonian migrants might have reached the Atlantic Forest during warmer temperature periods and consequently forest expansions. The timing of this colonization coincides with the establishment of forested pathways connecting southeastern Amazonia and the central-north Atlantic Forest (Batalha-Filho et al., 2013; Costa, 2003; Pavan et al., 2016).

Our empirical data help to reconstruct a more holistic picture of the South American rainforest history. While estimating speciation in Neotropical tree squirrels, we corroborate recent findings suggesting that much of the extant species-level diversity in Amazonia is young, dating back to the Quaternary (see Cracraft et al., 2020). The young speciation dates inferred for Neotropical tree squirrels indicated that their diversification likely occurred after the formation of the main Amazonian drainages (Hoorn et al., 2010). Therefore, it is likely that the Amazonian rivers acted mostly as barrier to secondary contact and gene flow. In other words, once a tree squirrel species crossed a river, its population remained isolated. This also leads us to infer that dispersal over vicariance was the dominant model of speciation in tree squirrels. A similar scenario of diversification was observed for one species of diurnal, tree-trunk specialist lizard (*Gonatodes humeralis*) and a species of tree frog (*Dendropsophus leucophyllatus*; Pirani et al., 2019). Besides the clear role of the riverine barriers in the migration and diversification of *Guerlinguetus* along the Amazonas basin, we suggest that climate-driven forest dynamics played important roles in the diversification, dispersion, and colonization of new areas by tree squirrels, particularly in the forests on the Atlantic coast of Brazil. Therefore, the current species diversity and the pattern of spatial organization of *Guerlinguetus* species must be a result of integrated effects from riverine barriers and climate-driven forest dynamics during the Pleistocene.

## Supporting information

Full description of Data Availability

Full description of the Supplementary Material

## Acknowledgments

We are deeply in debt to collectors, curators, and collection support staff, who provided us with preserved tissue samples or allowed us to obtain destructive samples from valuable museum specimens under their care: R.S. Voss, E. Hoeger, and N. Duncan (AMNH); M.R. Alvarez (CMARF); B.D. Patterson and A.W. Ferguson (FMNH); C.R. Silva and A.F. Sobrinho (IEPA); F. Catzeflis (ISEM); E. Pasa (MCN-FZB); C.G. Costa (MCN-M); A.U. Christoff, F.B. Peters and A.M. Gandini (MCNU); J.A. Oliveira and M. Weksler (MN); J. Silva Junior (MPEG); M.T. Rodrigues and B.M.S. Batista (MTR); C. Conroy, J.L. Patton, and F.J.M. Pascal (MVZ); M. Vivo, L.F. Silveira and J.G. Barros (MZUSP); L.P. Costa, Y.L.R. Leite and M.P. Nascimento (UFES-CTA); A.C. Mendes Oliveira (UFPA); J. Cherem, M. Graipel and E.C. Grisard (UFSC); D.P. Lunde, S.C. Peurach, N.R. Edmison, I. Rochon, M. Krol, J. Jacobs, C. Ludwig, C. Huddleston, D. DiMichele, M. Braun, L.H. Emmons, and A.L. Gardner (USNM). A. Ravetta, G.T. Garbino and L.P. Godoy provided tissue samples of recently collected specimens. We thank B. McLean and E. Schultz, who shared scripts and insights on introgression and migration analyses, respectively. We are also thankful to F. Burbrink, S. Castañeda-Rico, L. Parker for helpful discussion on analytical approaches and results interpretation. We are grateful to R. Dikow for providing technical support with the Smithsonian HPC. We finally thank L. Tomazelli and P. Mascaretti, who allowed us to use their photographs of *G. aestuans* and *G. brasiliensis*, respectively, depicted on Figure 3.

## Funding

This study had financial support from the Brazilian “Conselho Nacional de Desenvolvimento Científico e Tecnológico” through doctoral fellowships to EFA (147145/2016-3, 203692/2017-9, and 165553/2017-0) and through a postdoctoral fellowship to SEP (302204/2020-2); from the Smithsonian Institution through postdoctoral fellowship to SEP and through research funds provided to SEP and JEM; and from the Richard Gilder Graduate School of the American Museum of Natural History through a postdoctoral fellowship (Gerstner Scholar) and research funds provided to EFA.

## Data Availability

Full description of Data Availability for this manuscript is presented in a separate file.

## Conflict of Interest

The authors declare no conflict of interest.

## Author Contributions

**Edson F. Abreu**: conceptualization, methodology, formal analysis, investigation, data curation, writing (original draft), visualization, project administration, funding acquisition; **Joyce R. Prado**: conceptualization, methodology, formal analysis, investigation, data curation, writing (original draft), visualization; **Jesús E. Maldonado**: conceptualization, writing (review and editing), resources, funding acquisition; **Don E. Wilson**: writing (review and editing), resources, funding acquisition; **Alexandre R. Percequillo**: conceptualization, writing (review and editing), resources; **Silvia E. Pavan**: conceptualization, writing (review & editing), project administration, funding acquisition.

## Notes

### Competing Interest Statement

The authors have declared no competing interest.

