## Supplementary material for "Quantitative species delimitation, tests for gene flow, and migration models uncover complex, recent speciation in tree squirrels": Full description of Data Availability

**Data Availability for the Manuscript:**

### Data Availability

#### Raw data

Raw sequence data (Illumina cleaned reads) for all 72 vouchered museum samples analyzed in this study (67 *Guerlinguetus* in addition to five outgroups) is publicly available through the National Center for Biotechnology Information (NCBI) Sequence Read Archive (SRA) under the BioProject PRJNA847638.

#### Data sets

All data sets produced in this study are publicly available as follow:

- **Data set 1**: UCE loci data set including 3,642 loci (1,495,138 bp) from 67 individuals of *Guerlinguetus* in addition to five outgroups. Available as PHYLIP file here: [XXX]
- **Data set 2**: 1,169 biallelic, unlinked SNPs called from 64 samples of *Guerlinguetus*. Available as a VCF file here: [XXX]
- **Data set 3**: Reduced UCE loci data set including a subset of 500 loci (corresponding to the loci with the highest sample representativeness) from 67 specimens of *Guerlinguetus* in addition to one outgroup. Available as NEXUS file here: [XXX]
- **Data set 4**: Reduced SNP data set including 1,155 SNPs from 17 specimens of *Guerlinguetus*. Available as a NEXUS file here: [XXX]

#### Codes

An entire GitHub repository including scripts, input files, and output files needed to fully reproduce the analyses performed in this manuscript is publicly available here: [XXX]
