## Supplementary material for "Quantitative species delimitation, tests for gene flow, and migration models uncover complex, recent speciation in tree squirrels": Full description of the Supplementary Material

#### Supplementary Table S1

Complete list of *Guerlinguetus* samples analyzed in this study, accompanying museum catalogue and geographic data.

| Voucher | Original species ID | Country | State/ Depart. | Municipality/ Locality | Loc. # | Latitude | Longitude |
|---|---|---|---|---|---|---|---|
| IEPA 3704 | *G. aestuans* | Brazil | Amapá | Porto Grande, margem direita do Rio Vila Nova, Floresta Estadual do Amapá | 1 | 0.470000 | -52.010000 |
| MN 56819 | *G. aestuans* | Brazil | Amazonas | Barcelos, Rio Katana-u | 2 | 1.208611 | -64.789167 |
| LMUSP (ENM 13) | *G. aestuans* | Brazil | Amazonas | São Gabriel da Cachoeira, 5º PEF Maturaca | 3 | 0.634605 | -66.124988 |
| CMARF 1568 | *G. brasiliensis* | Brazil | Bahia | Belmonte, Fazenda Ouro Verde | 4 | -15.895070 | -39.238440 |
| CMARF 1569 | *G. brasiliensis* | Brazil | Bahia | Belmonte, Fazenda Ouro Verde | 4 | -15.895070 | -39.238440 |
| UFES-CTA 85 | *G. brasiliensis* | Brazil | Bahia | Nova Viçosa, Fazenda Suécia | 5 | -17.878889 | -40.026389 |
| UFES-CTA 223 | *G. brasiliensis* | Brazil | Espírito Santo | Águia Branca, Fazenda Pedra Redonda | 6 | -18.973333 | -40.770556 |
| UFES-CTA 3817 | *G. brasiliensis* | Brazil | Espírito Santo | Conceição da Barra, Floresta Nacional do Rio Preto | 7 | -18.355278 | -39.844167 |
| UFES-CTA 119 | *G. brasiliensis* | Brazil | Espírito Santo | Pancas, Córrego São Bento, Fazenda do Dr. Rolly Luís | 8 | -19.225833 | -40.761667 |
| UFES-CTA 614 | *G. brasiliensis* | Brazil | Espírito Santo | Viana, Pimenta | 9 | -20.379167 | -40.468333 |
| UFES-CTA 14 | *G. brasiliensis* | Brazil | Espírito Santo | Vitória, Parque Estadual da Fonte Grande | 10 | -20.350000 | -40.350000 |
| MPEG (DICO 001) | *G. aestuans* | Brazil | Mato Grosso | Reserva Extrativista Guariba-Roosevelt, margem direita Rio Roosevelt | 11 | -9.000556 | -60.354167 |
| UFES-CTA 3776 | *G. brasiliensis* | Brazil | Minas Gerais | Belo Horizonte, Parque das Mangabeiras | 12 | -19.945590 | -43.910533 |
| MCN-M 866 | *G. brasiliensis* | Brazil | Minas Gerais | Braúna, UHE Porto Estrela | 13 | -19.116389 | -42.657778 |
| MCN-M 2944 | *G. brasiliensis* | Brazil | Minas Gerais | Brumadinho, Mina do Córrego do Feijão | 14 | -20.113931 | -44.115419 |
| UFES-CTA 1063 | *G. brasiliensis* | Brazil | Minas Gerais | Itanhandu, Posses, 13 km SE Itanhandu | 15 | -22.383333 | -44.850000 |
| MPEG (CN 158) | *G. aestuans* | Brazil | Pará | Alenquer, ESEC Grão-Pará, porção sul | 16 | -0.165489 | -55.186400 |
| MPEG (CN 160) | *G. aestuans* | Brazil | Pará | Alenquer, ESEC Grão-Pará, porção sul | 16 | -0.165489 | -55.186400 |
| IEPA 4384 | *G. aestuans* | Brazil | Pará | Almeirim, margem direita do Rio Jari, Igarapé Pacanari | 17 | 0.681910 | -52.593170 |
| MPEG (CN 269) | *G. aestuans* | Brazil | Pará | Almerim, Flota Paru, margem direita do rio Paru de Leste | 18 | -0.943969 | -53.236300 |

| | | | | | | | |
|---|---|---|---|---|---|---|---|
| MPEG (CN 270) | *G. aestuans* | Brazil | Pará | Almerim, Flota Paru, margem direita do rio Paru de Leste | 18 | -0.943969 | -53.236300 |
| MPEG (CN 230) | *G. aestuans* | Brazil | Pará | Almerim, Reserva Biológica Maicuru | 19 | -0.828619 | -53.931200 |
| MPEG (CN 241) | *G. aestuans* | Brazil | Pará | Almerim, Reserva Biológica Maicuru | 19 | -0.828619 | -53.931200 |
| UFMT (BML 1493) | *G. aestuans* | Brazil | Pará | Belo Monte | 20 | -3.400000 | -51.750000 |
| MPEG (CN 029) | *G. aestuans* | Brazil | Pará | Faro, Flota de Faro, margem esquerda do Rio Nhamundá | 21 | -1.714011 | -57.213300 |
| MPEG (CN 048) | *G. aestuans* | Brazil | Pará | Faro, Flota de Faro, margem esquerda do Rio Nhamundá | 21 | -1.714011 | -57.213300 |
| MPEG (PECC 26) | *G. aestuans* | Brazil | Pará | Marajó-Afuá, Igarapé Torrão, rio Preto, PE Charapucú | 22 | -0.418840 | -50.494876 |
| MPEG (PECC 03) | *G. aestuans* | Brazil | Pará | Marajó-Afuá, Rio Cuieiras, PE Charapucú | 23 | -0.226960 | -50.590000 |
| MPEG (CN 294) | *G. aestuans* | Brazil | Pará | Óbidos, ESEC Grão-Pará, porção central | 24 | -0.630281 | -55.728500 |
| MPEG (CN 175) | *G. aestuans* | Brazil | Pará | Oriximiná, ESEC Grão-Pará, porção norte | 25 | -1.285419 | -58.695900 |
| MPEG (CN 201) | *G. aestuans* | Brazil | Pará | Oriximiná, ESEC Grão-Pará, porção norte | 25 | -1.285419 | -58.695900 |
| LMUSP (EFA 41) | *G. brasiliensis* | Brazil | Pará | Pacajá, LT Xingu-Estreito | 26 | -3.929722 | -51.069722 |
| MCN-M 1388 | *G. brasiliensis* | Brazil | Pará | Parauapebas, FLONA de Carajás | 27 | -6.220806 | -50.298406 |
| MPEG (ANRA 10) | *G. brasiliensis* | Brazil | Pará | Portel, Igarapé Açaituba | 28 | -2.224211 | -50.544875 |
| MPEG (ANRA 12) | *G. brasiliensis* | Brazil | Pará | Portel, Igarapé Quirino | 29 | -2.166682 | -50.644508 |
| USNM 549526 | *G. brasiliensis* | Brazil | Pará | Rio Xingu, east bank | 30 | -3.650000 | -52.370000 |
| MZUSP (GTG 54) | *G. brasiliensis* | Brazil | Pará | Santana do Araguaia, Fazenda Fartura | 31 | -9.732500 | -50.325278 |
| MZUSP (GTG 55) | *G. brasiliensis* | Brazil | Pará | Santana do Araguaia, Fazenda Fartura | 31 | -9.732500 | -50.325278 |
| MPEG 45441 | *G. aestuans* | Brazil | Pará | Santarém, Comunidade Alto Mentai | 32 | -2.800250 | -55.583570 |
| MPEG (RETA 15) | *G. aestuans* | Brazil | Pará | Santarém, Comunidade de Capixauã | 33 | -2.612361 | -55.192083 |
| UFPA (JMIJ 35) | *G. aestuans* | Brazil | Pará | Tapajós | 34 | -5.613410 | -57.122430 |
| LMUSP (BM 10678) | *G. aestuans* | Brazil | Pará | Vitória do Xingu, margem esquerda do Rio Xingu | 35 | -2.873760 | -52.015519 |
| LMUSP (BM 17058) | *G. aestuans* | Brazil | Pará | Vitória do Xingu, margem esquerda do Rio Xingu | 35 | -2.873760 | -52.015519 |
| UFES-CTA 4256 | *G. aestuans* | Brazil | Pará | Vitória do Xingu, margem esquerda do Rio Xingu | 36 | -3.259346 | -51.856208 |
| MTR-CIT 1128 | *G. brasiliensis* | Brazil | Rio Grande do Sul | Itá | 37 | -27.256812 | -52.392234 |
| FZB 3514 | *G. brasiliensis* | Brazil | Rio Grande do Sul | Machadinho | 38 | -27.565923 | -51.659475 |

| | | | | | | | |
|---|---|---|---|---|---|---|---|
| MCNU 3683 | *G. brasiliensis* | Brazil | Santa Catarina | Anita Garibaldi, margem direita UHE Barra Grande | 39 | -27.788560 | -51.154506 |
| UFSC 2930 | *G. brasiliensis* | Brazil | Santa Catarina | Ipuaçu, AHE Quebra Queixo | 40 | -26.657551 | -52.547408 |
| UFSC 4047 | *G. brasiliensis* | Brazil | Santa Catarina | Xaxim, Arvoredo, PCH Arvoredo | 41 | -27.040977 | -52.466292 |
| FMNH 141601 | *G. brasiliensis* | Brazil | Sao Paulo | Ilha do Cardoso | 42 | -25.133333 | -47.966667 |
| MTR-CIT 1785 | *G. brasiliensis* | Brazil | São Paulo | Biritiba Mirim | 43 | -23.596572 | -46.026164 |
| MVZ 192702 | *G. brasiliensis* | Brazil | São Paulo | Capão Bonito, Base do Carmo, Fazenda Intervales | 44 | -24.333333 | -48.416667 |
| LMUSP (DTM 277) | *G. brasiliensis* | Brazil | São Paulo | Caraguatatuba, Parque Estadual da Serra do Mar | 45 | -23.581412 | -45.484681 |
| LMUSP (DTM 41) | *G. brasiliensis* | Brazil | São Paulo | Caraguatatuba, Parque Estadual da Serra do Mar | 45 | -23.581412 | -45.484681 |
| MTR-CIT 1821 | *G. brasiliensis* | Brazil | São Paulo | Juquitiba | 46 | -23.928496 | -47.066410 |
| MTR-ITM 444 | *G. brasiliensis* | Brazil | São Paulo | Juquitiba | 46 | -23.928496 | -47.066410 |
| MTR-ITM 192 | *G. brasiliensis* | Brazil | São Paulo | Piedade | 47 | -23.716279 | -47.422659 |
| MVZ 200389 | *G. brasiliensis* | Brazil | São Paulo | São Sebastião, Fazenda da Toca, 2.4 km E, 0.8 km NE (by road) Ilhabela, Ilha de São Sebastião | 48 | -23.816667 | -45.350000 |
| LMUSP 301 | *G. brasiliensis* | Brazil | São Paulo | Sorocaba, APP Toyota | 49 | -23.374444 | -47.470833 |
| MVZ 182070 | *G. brasiliensis* | Brazil | São Paulo | Ubatuba, Fazenda Capricórnio, 5 km N Ubatuba | 50 | -23.416667 | -45.116667 |
| MVZ 182071 | *G. brasiliensis* | Brazil | São Paulo | Ubatuba, Praia do Félix | 51 | -23.383333 | -44.466667 |
| ISEM-T 5176 | *G. aestuans* | French Guiana | Cacao | Roura, Cacao | 52 | 4.576944 | -52.468611 |
| ISEM-T 6101 | *G. aestuans* | French Guiana | Cayenne | Camp du Tigre, Cayenne | 53 | 4.908333 | -52.308333 |
| ISEM-T 1758 | *G. aestuans* | French Guiana | Petit Saut | Sinnamary, Petit Saut | 54 | 5.050000 | -53.050000 |
| USNM 599925 | *G. aestuans* | Guyana | | Mount Roraima, N slope | 55 | 5.283300 | -60.750000 |
| USNM 560645 | *G. aestuans* | Venezuela | Amazonas | Neblina Base Camp, Rio Mawarinuma | 56 | 0.830000 | -66.170000 |
| AMNH 135435 | *G. aestuans* | Venezuela | Bolívar | Gran Sabana, Camarata Valley | 57 | 5.500000 | -61.500000 |

**Supplementary Table S2**

The ΔK values for the Structure analyses based on mean likelihood (Mean LnP(K)) and standard deviation (Stdev LnP(K)) for K = 1–15. The ΔK values for the two most probable numbers of clusters according to the Evanno method are 18344.935190 (K = 2) and 9.50545 (K = 6).

| K | Mean LnP(K) | Stdev LnP(K) | Delta K |
|---|---|---|---|
| 1 | -56531.980000 | 2.580611 | — |
| 2 | -36498.030000 | 0.922617 | 18344.935190 |
| 3 | -33389.430000 | 949.943788 | 0.024054 |
| 4 | -30303.680000 | 756.762555 | 0.750301 |
| 5 | -27785.730000 | 259.369908 | 9.056910 |
| 6 | -27616.870000 | 2781.580912 | 9.505451 |
| 7 | -53888.190000 | 52668.572423 | 0.284003 |
| 8 | -65201.480000 | 42349.154402 | 1.056067 |
| 9 | -31791.210000 | 16981.601288 | 2.222667 |
| 10 | -36125.380000 | 31292.935037 | 0.177682 |
| 11 | -46019.730000 | 37081.848322 | 0.609209 |
| 12 | -33323.470000 | 8520.502071 | 1.212460 |
| 13 | -30957.980000 | 5814.712840 | 0.724184 |
| 14 | -32803.410000 | 6483.240940 | 0.038223 |
| 15 | -34401.030000 | 9832.667065 | — |

**Supplementary Table S3**

Statistics calculated by Dsuite (Patterson's D, Z-score, and f4-ratio) for all possible species trios with a pre-determined fourth species (Tepuis) used as outgroup.

| P1 | P2 | P3 | D-statistic | Z-score | p-value | f4-ratio | BBAA | ABBA | BABA |
|---|---|---|---|---|---|---|---|---|---|
| Atlantic Forest | Xingu-Tocantins | Madeira-Tapajós | 0.065 | 0.759 | 0.448 | 0.018 | 152.824 | 11.303 | 9.933 |
| Xingu-Tocantins | Northern Amazonia | Madeira-Tapajós | 0.052 | 0.564 | 0.573 | 0.034 | 39.401 | 26.069 | 23.505 |
| Xingu-Tocantins | Eastern Amazonia | Madeira-Tapajós | 0.059 | 0.858 | 0.391 | 0.034 | 61.439 | 23.475 | 20.853 |
| Atlantic Forest | Northern Amazonia | Madeira-Tapajós | 0.077 | 0.899 | 0.368 | 0.051 | 37.433 | 27.388 | 23.454 |
| Atlantic Forest | Eastern Amazonia | Madeira-Tapajós | 0.085 | 1.094 | 0.274 | 0.052 | 58.670 | 25.455 | 21.462 |
| Northern Amazonia | Eastern Amazonia | Madeira-Tapajós | 0.001 | 0.012 | 0.991 | 0.001 | 41.477 | 28.825 | 28.767 |
| Atlantic Forest | Xingu-Tocantins | Northern Amazonia | 0.131 | 1.737 | 0.082 | 0.059 | 139.856 | 14.231 | 10.944 |
| Atlantic Forest | Xingu-Tocantins | Eastern Amazonia | 0.172 | 2.300 | 0.021 | 0.110 | 117.226 | 16.352 | 11.542 |
| Eastern Amazonia | Xingu-Tocantins | Northern Amazonia | 0.011 | 0.146 | 0.884 | 0.012 | 53.170 | 28.419 | 27.797 |
| Atlantic Forest | Eastern Amazonia | Northern Amazonia | 0.045 | 0.476 | 0.634 | 0.048 | 51.216 | 30.653 | 27.988 |

**Supplementary Table S4**

Results for demographic parameters (mean and 95% high posterior densities) estimated using G-PhoCs. Effective population size (Ne) and Divergence time (Tdiv) are shown.

| Demographic Parameters | Mean | 95% HPD lower | 95% HPD upper |
|---|---|---|---|
| Ne Tepuis | 115,750 | 98,333 | 131,667 |
| Ne Madeira-Tapajós | 93,808.33 | 80,833.33 | 104,166.67 |
| Ne Northern-Amazonia | 143,108.33 | 129,166.67 | 156,666.67 |
| Ne Eastern-Amazonia | 209,775 | 191,666.67 | 228,333.33 |
| Ne Xingu-Tocantins | 125,891.67 | 114,166.67 | 140,000 |
| Ne Atlantic Forest | 92,225 | 83,333.33 | 100,000 |
| Tdiv root | 1,878,375 | 1,740,000 | 2,002,500 |
| Tdiv Atlantic Forest/ Xingu-Tocantins/ Eastern-Amazonia/ Northern-Amazonia/ Madeira-Tapajós | 1,148,550 | 1,072,500 | 1,215,000 |
| Tdiv Atlantic Forest/ Xingu-Tocantins/ Eastern-Amazonia/ Northern-Amazonia | 458,025 | 412,500 | 495,000 |
| Tdiv Atlantic Forest/ Xingu-Tocantins/ Eastern-Amazonia | 447,863 | 412,500 | 487,500 |
| Tdiv Atlantic Forest/ Xingu-Tocantins | 424,575 | 390,000 | 457,500 |

**Supplementary Information S1**

Bioinformatics pipeline used to phase UCE data, call, and filter SNPs.

---

**Per locus trimmed sequence alignments**

(1) Follow the PHYLUCE 1.6 pipeline (Faircloth, 2016) up to the step of "Alignment cleaning" to generate individual (per locus) alignments for all UCEs enriched. Alignments will be saved in NEXUS format.

**Pseudogenomic reference**

(1) Import to Geneious all per locus NEXUS alignments;
(2) Generated consensus sequences for each locus in Geneious;
(3) Concatenated all sequences to build a pseudogenomic reference in Geneious;
(4) Export the pseudogenomic reference in FASTA format;
(5) Create an .dic file for the pseudogenomic reference using GATK 4.2.1.0:

```
rungatk CreateSequenceDictionary -R ref.fasta
```

(6) Create an .fai file for the pseudogenomic reference using SAMtools 1.14:

```
samtools faidx ref.fasta
```

**Reference sample**

(1) Select the sample with the greatest number of UCEs enriched to be the reference sample;
(2) Create an .dic file for the reference sequence using GATK 4.2.1.0:

```
rungatk CreateSequenceDictionary -R ref.fasta
```

(3) Create an .fai file for the reference sequence using SAMtools 1.14:

```
samtools faidx ref.fasta
```

**BAM files**

(1) Use BWA-mem algorithm implemented through PHYLUCE to map cleaned FASTQ reads for each sample against the pseudogenomic reference or the reference sequence. This will result in indexed BAM files for each sample:

```
phyluce_snp_bwa_multiple_align --config phasing.conf \
--output multialign-bams \
--log-path log \
--mem
```

**Mark duplicates**

(1) Remove duplicates reads using Picard:

```
java -jar picard.jar MarkDuplicates I=[sample].bam
O=marked_duplicates.bam M=marked_dup_metrics.txt
```

**Call SNPs and Indels separately for each sample**

(1) Use GATK to create a gVCF file for each sample:

```
rungatk HaplotypeCaller \
-R reference.fasta \
-I [sample]-CL-RG-MD-M.bam \
-O [sample].g.vcf.gz \
-ERC GVCF
```

**Merge and genotype gVCF files**

(1) Unite all gVCF files into a single folder;
(2) Run GATK to merge all of the different gVCF files into one file:

```
rungatk CombineGVCFs \
-R reference.fasta \
-V [sample1].g.vcf.gz -V [sample2].g.vcf.gz ...
-O all-gatk-combine.g.vcf.gz
```

(3) Run GATK to genotype all sample:

```
rungatk GenotypeGVCFs \
-R reference.fasta \
-V all-gatk-combine.g.vcf.gz \
-O all-gatk-genotype.g.vcf.gz
```

**Extract SNPs from the call set**

(1) Run GATK to extract both SNPs and Indels from the call set:

```
rungatk SelectVariants \
-R reference.fasta \
-V all-gatk-genotype.g.vcf.gz \
--select-type-to-include SNP \
-O all-gatk-select-variants.g.vcf.gz
```

**Variant Filtering**

(1) Run GATK to filter the SNPs to remove low-quality calls:

```
rungatk VariantFiltration \
-R reference.fasta \
-V all-gatk-select-variants.g.vcf.gz \
-O all-gatk-filter-variants.g.vcf.gz
--filterExpression "QD < 2.0 || FS > 60.0 || MQ < 40.0 ||
HaplotypeScore > 13.0 || MappingQualityRankSum < -12.5 ||
ReadPosRankSum < -8.0" \
--filterName "my_snp_filter"
```

(2) Generate a VCF file using VCFtools:

```
vcftools --gzvcf all-gatk-filter-variants.g.vcf.gz --remove-
filtered-all --maf 0.05 --minDP 2 --recode --out all-spns.vcf
```

(3) Remove filtered sites from the VCF file and selected SNPs with a minimum depth of coverage of 5 per individual and a minor allele frequency ≥ 5% using VCFtools:

```
vcftools --vcf all-spns.vcf.recode.vcf --remove-filtered-all
--maf 0.05 --mac 5 --recode --out thinned_snps.vcf
```

(4) Use HD_plot.R to filter SNPs resulting from putative paralogues or incorrectly assembled contigs from the data set by removing SNPs with heterozygosity >0.75 and a read-ratio deviation score D >10:

```
Het&D.R # save file with SNPs to exclude

bcftools annotate --set-id +'%CHROM\:%POS' thinned_snps.vcf >
thinned_snps_edited.vcf # edit vcf ID column

vcftools --vcf thinned_snps_edited.vcf --exclude
snp_to_exclude.txt --recode --out excluded_snps.vcf
```

(5) Select unliked SNPs:

```
vcftools --vcf excluded_snps.vcf --thin 2000 --recode --out
unlinked_snps.vcf
```

(6) Filter SNPs by missing data:

```
vcftools --vcf unlinked_snps.vcf --missing-indv --out sub-
missind
```

(7) Remove samples with great amount of missing data:

```
vcftools --vcf unlinked_snps.vcf --remove-indv IND_XX --
recode --out unlinked_excluded_ind.vcf
```

(8) Remove loci with great amount of missing data:

```
vcftools --vcf unlinked_excluded_ind.vcf --max-missing 0.75 -
-recode --out final_25%_missing_snps.vcf
```

---

#### Supplementary Information S2

Geographic ranges of the population clusters inferred by the DAPC and hierarchical Structure (K = 9):

**Tepuis**: includes samples from the flat top mountains on the Guiana Shield.

**Madeira-Tapajós**: includes sampled from the southern bank of Solimões River, on the interfluve of Madeira and Tapajós rivers.

**Northern Amazonia (west)**: includes samples from the northern bank of Amazonas River, on the interfluve of Negro and Amazonas rivers.

**Northern Amazonia (east)**: includes samples from the northern bank of Amazonas River, from the right bank of Negro River eastwards to Jari River basin.

**Eastern Amazonia (north)**: includes samples from the eastern portion of the northern bank of Amazonas River, east of the Jari River.

**Eastern Amazonia (south)**: includes individuals from the interfluve of Tapajós and Xingu rivers, on the southern bank of Amazonas River.

**Xingu-Tocantins**: includes samples from southeastern Amazonia, on the interfluve of Xingu and Tocantins Rivers, on the southern bank of Amazonas River.

**Atlantic Forest (north)**: includes samples from the Jequitinhonha River to the northern bank on the headwaters of Tietê River.

**Atlantic Forest (south)**: includes samples from the Tietê River southwards to Rio Uruguay.

#### Supplementary Figure S1

Collecting localities of all specimens of *Guerlinguetus* analyzed in this study. Symbols are colored according to the proposed species hypothesis: Tepuis (brown), Madeira-Tapajós (peach), Northern Amazonia (light green), Eastern Amazonia (navy blue), Xingu-Tocantins (pink), Atlantic Forest (purple). Localities numbers follow Supplementary Table S1.

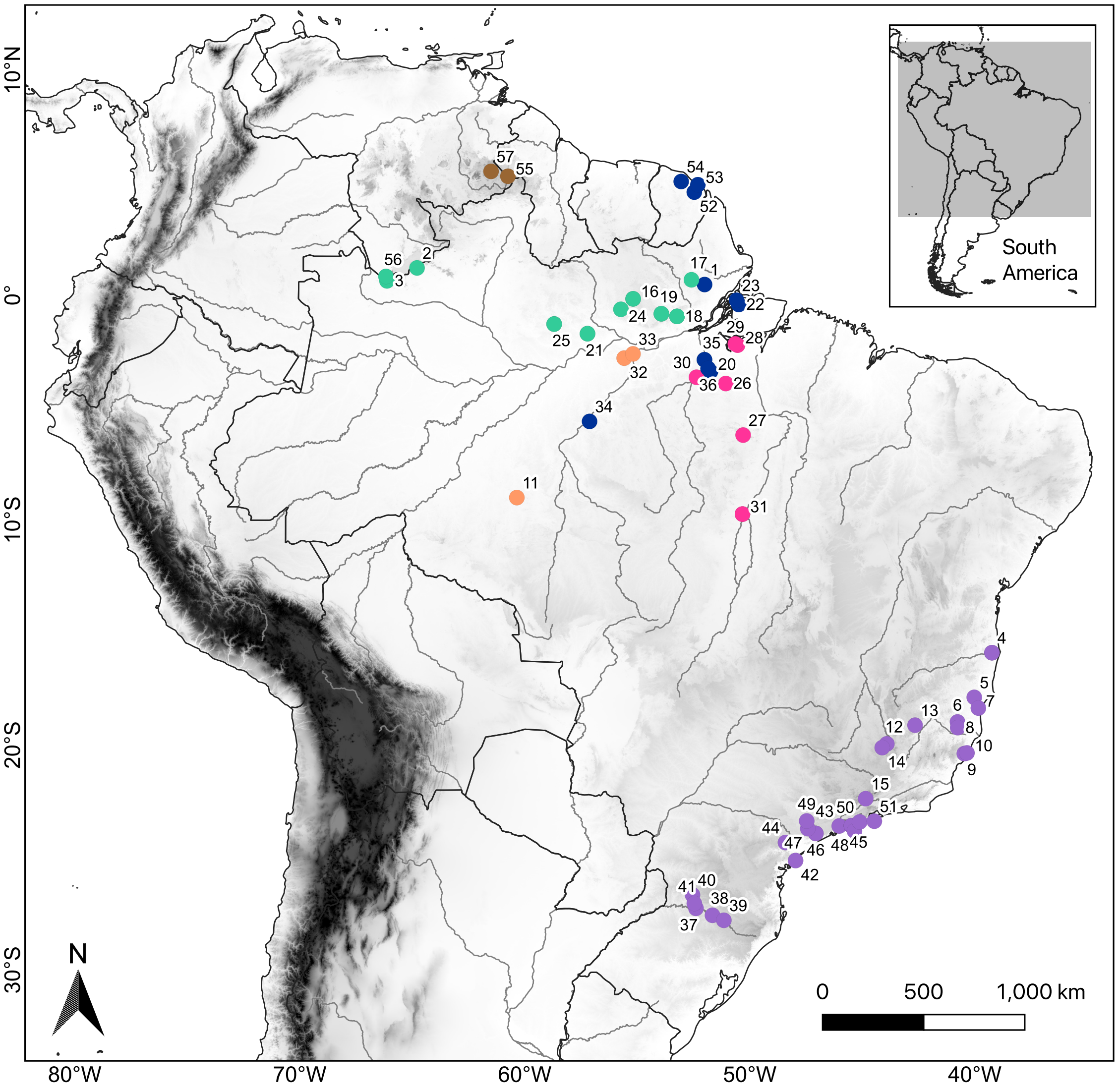

**Supplementary Figure S2**

Comparison on the clustering results (1–15) from the DAPC based on the Bayesian Information Criterion (BIC) showing K = 9 as the lowest value.

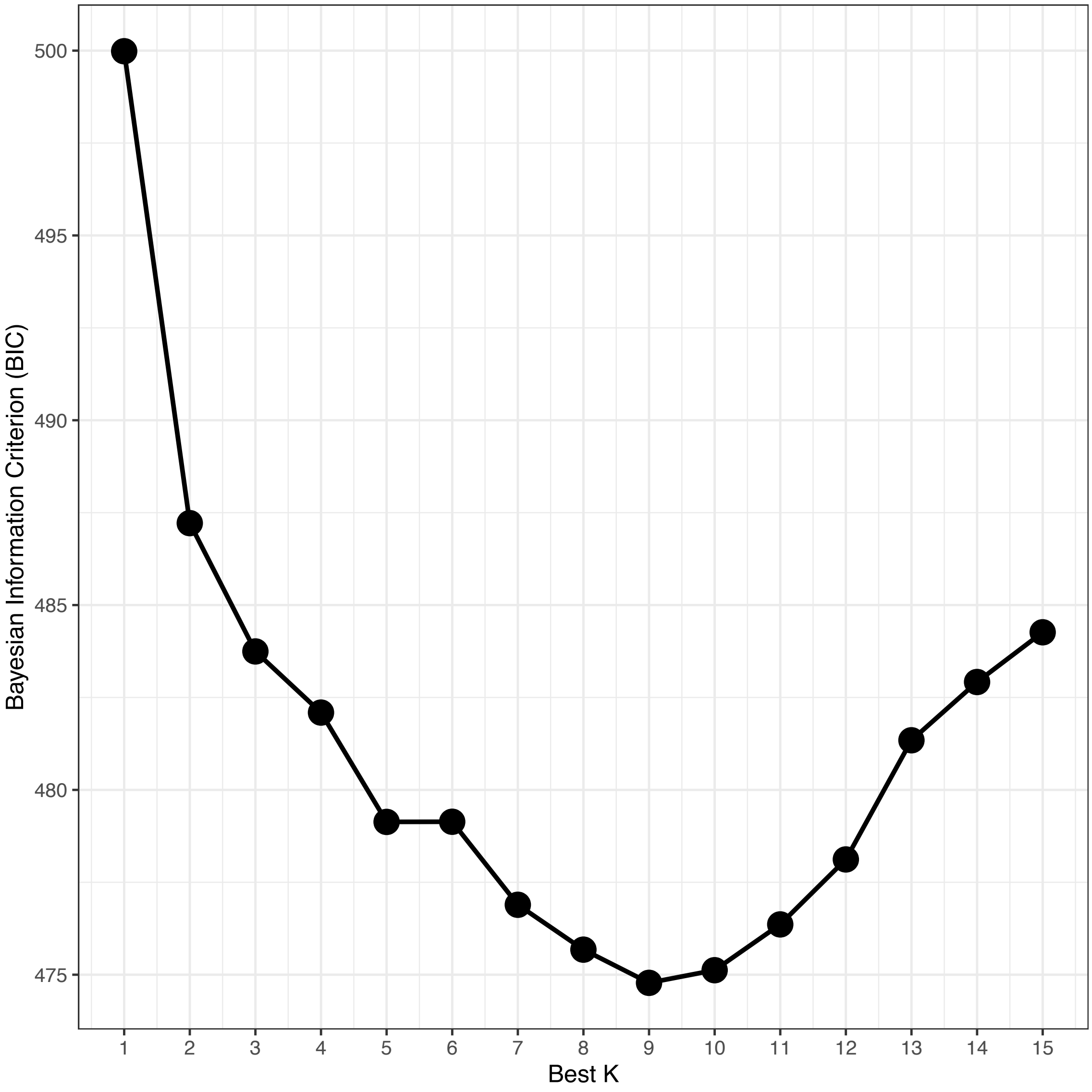

**Supplementary Figure S3**

The ΔK values for the Structure analyses based on mean likelihood (Mean LnP(K)) and standard deviation (Stdev LnP(K)) for K = 1–15. The ΔK values for the two most probable numbers of clusters according to the Evanno method are 18344.935190 (K = 2) and 9.50545 (K = 6).

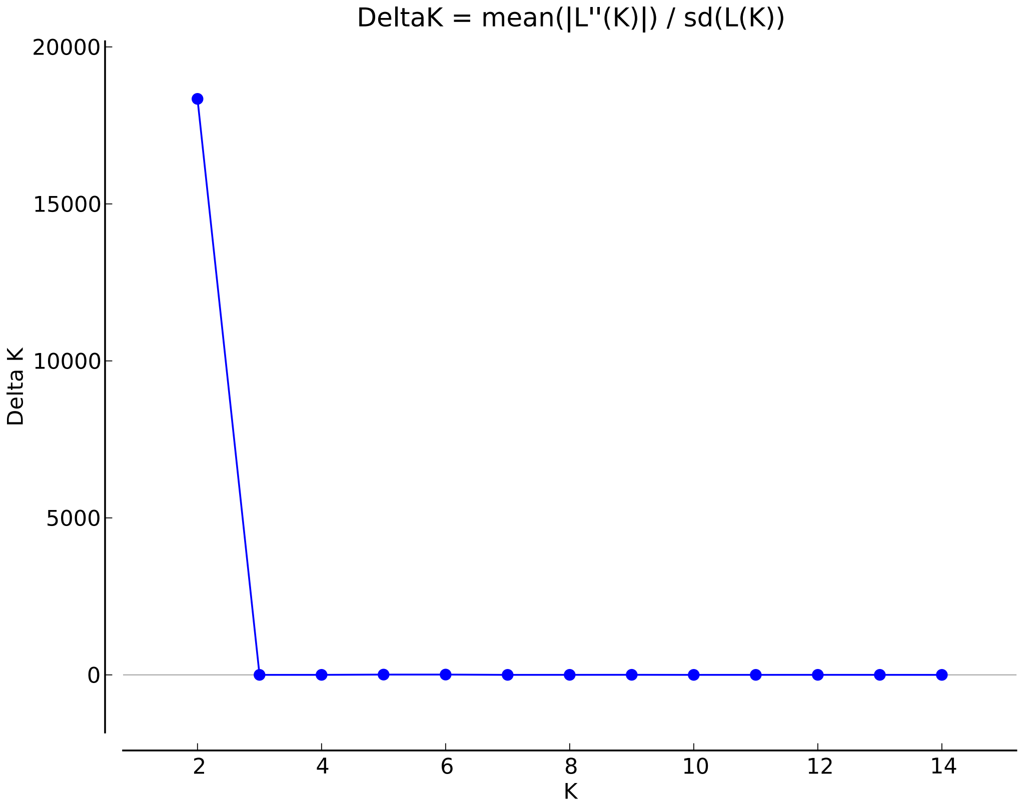

#### Supplementary Figure S4

Population structure and ancestry assignments of *Guerlinguetus* based on the hierarchical Structure analysis of SNP data (1,169 SNPs, 64 specimens). a) Phylogenetic relationships as inferred from the coalescent approach in SVDquartets, with correspondence to the bar plots. b) Ancestry assignments and genetic admixture shown as bar plots for each individual after three rounds of analyses. c) Pie charts showing ancestry assignments and genetic admixture for each individual after three rounds of analyses (K = 9) plotted on the map according to the sampling localities. The three terminals of the phylogeny colored in black are specimens exclusively analyzed for UCEs but not for SNPs.

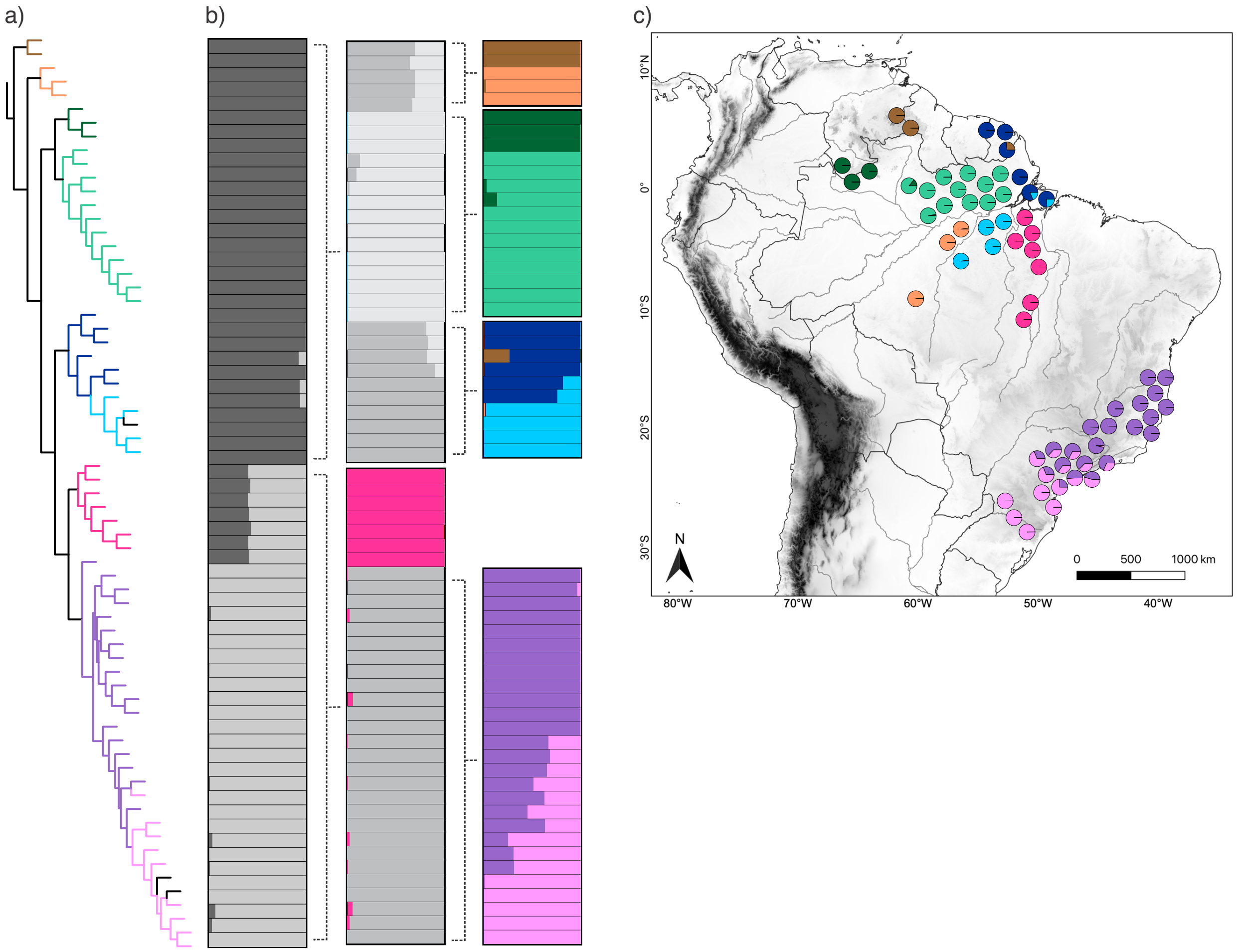

**Supplementary Figure S5**

Mantel test result showing correlation between geographical distance (km) and genetic distance ($F_{st}$) for Atlantic Forest populations.

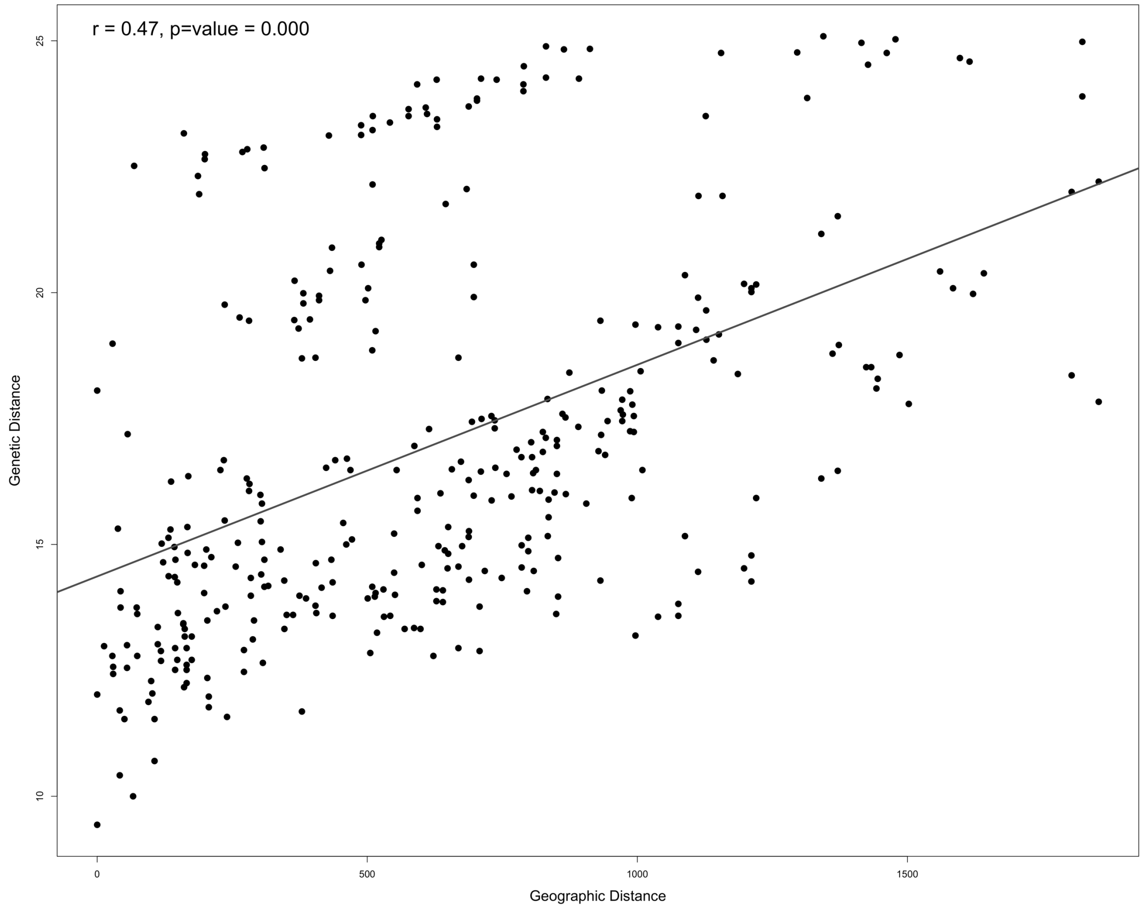

**Supplementary Figure S6**

Mantel test result showing correlation between geographical distance (km) and genetic distance ($F_{st}$) for Northern Amazonian populations.

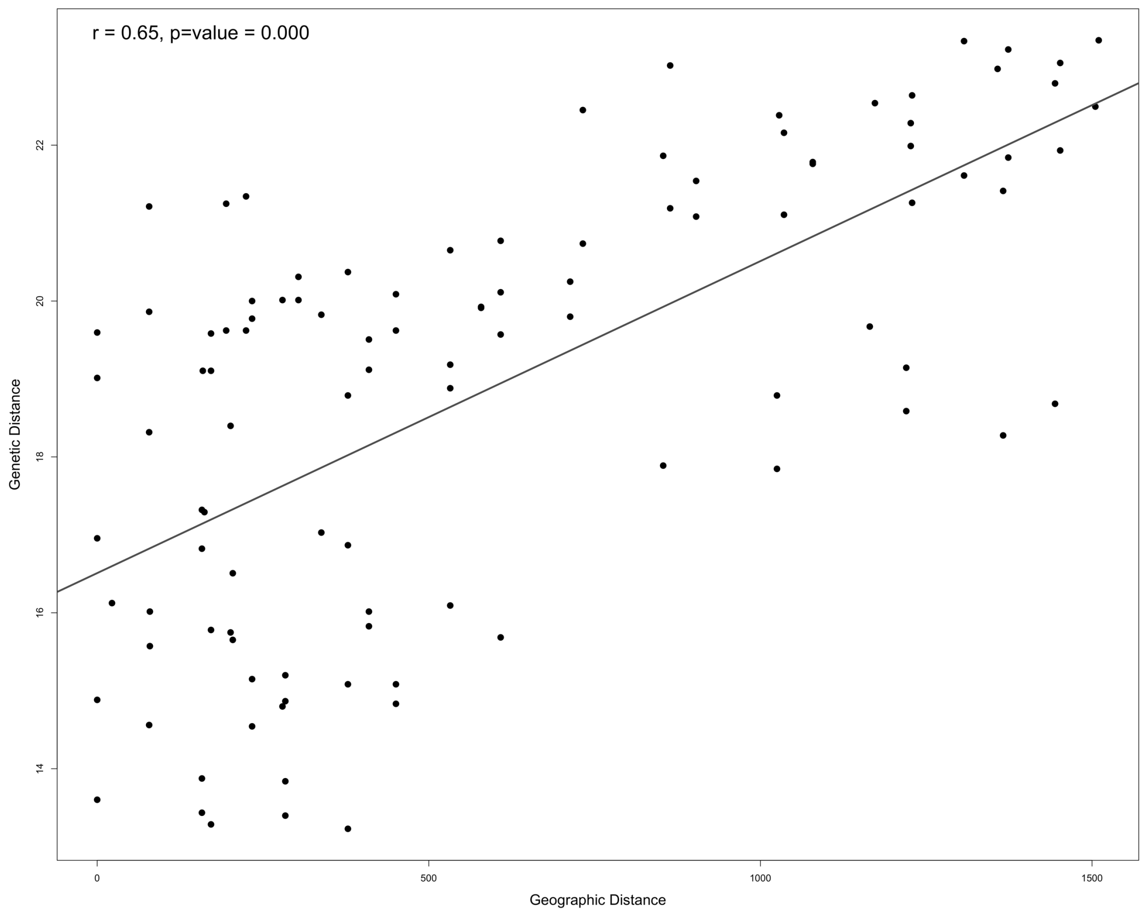

#### Supplementary Figure S7

Mantel test result showing correlation between geographical distance (km) and genetic distance ($F_{st}$) for Eastern Amazonian populations.

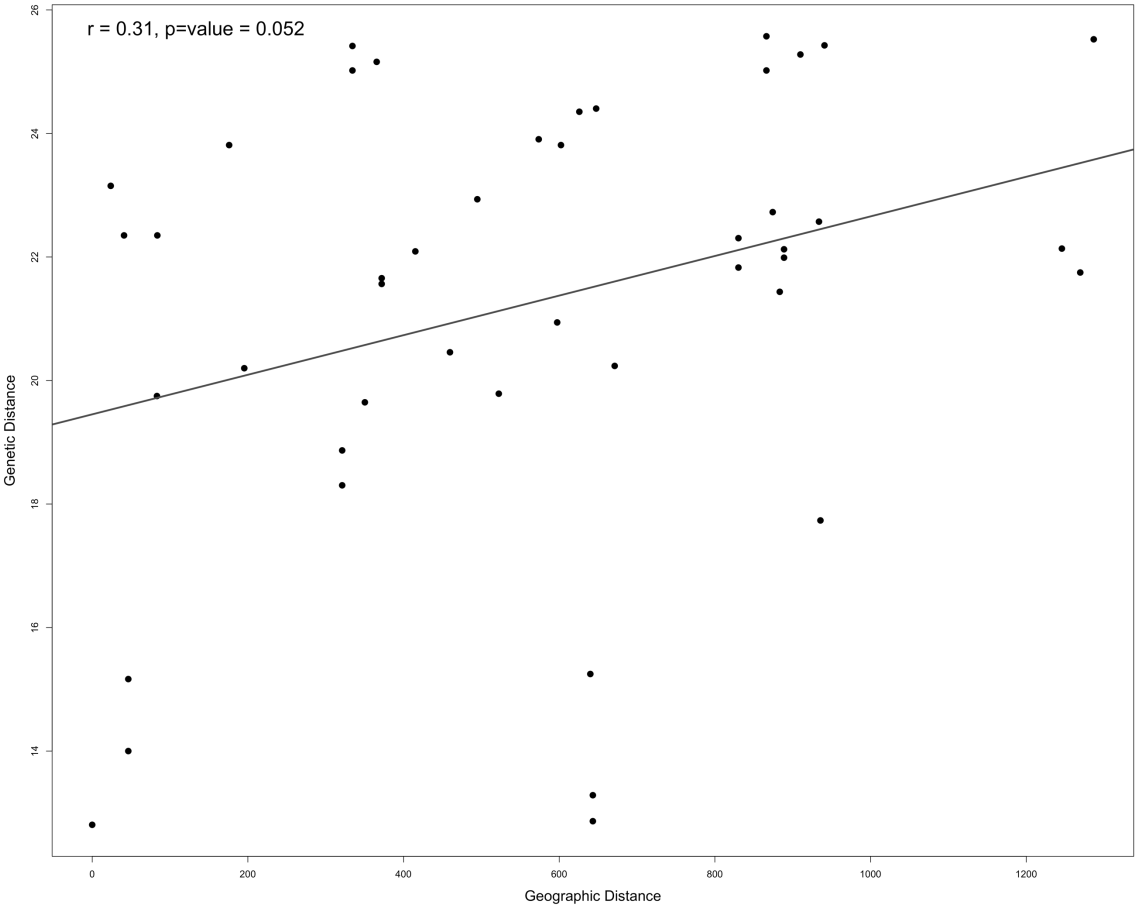

### Supplementary Figure S8

Phylogenetic reconstruction based on loci (data set including 3,642 UCEs and 67 individuals of *Guerlinguetus*) inferred with RAxML.

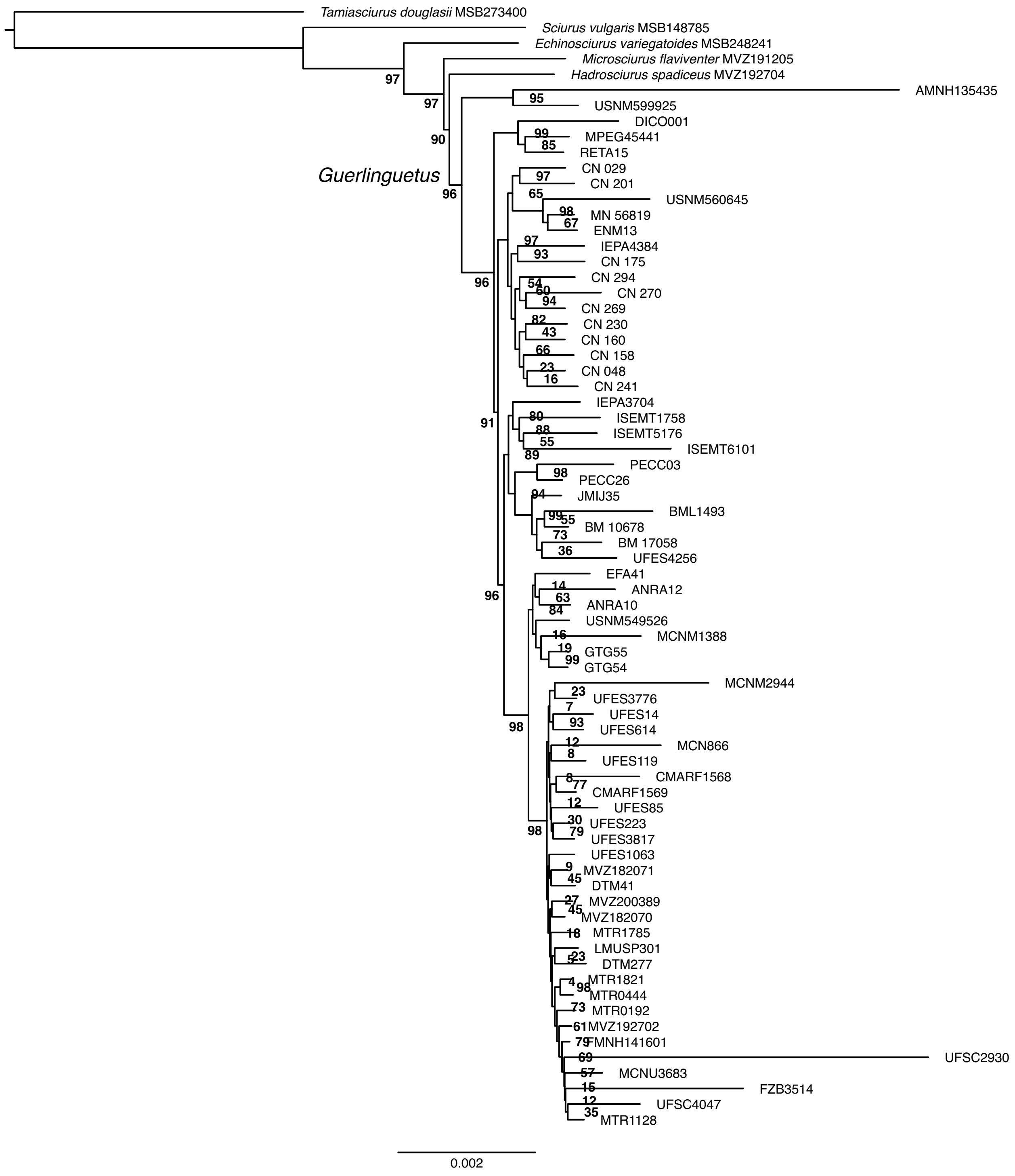

### Supplementary Figure S9

Phylogenetic reconstruction based on loci (data set including 3,642 UCEs and 67 individuals of *Guerlinguetus*) inferred with IQ-Tree.

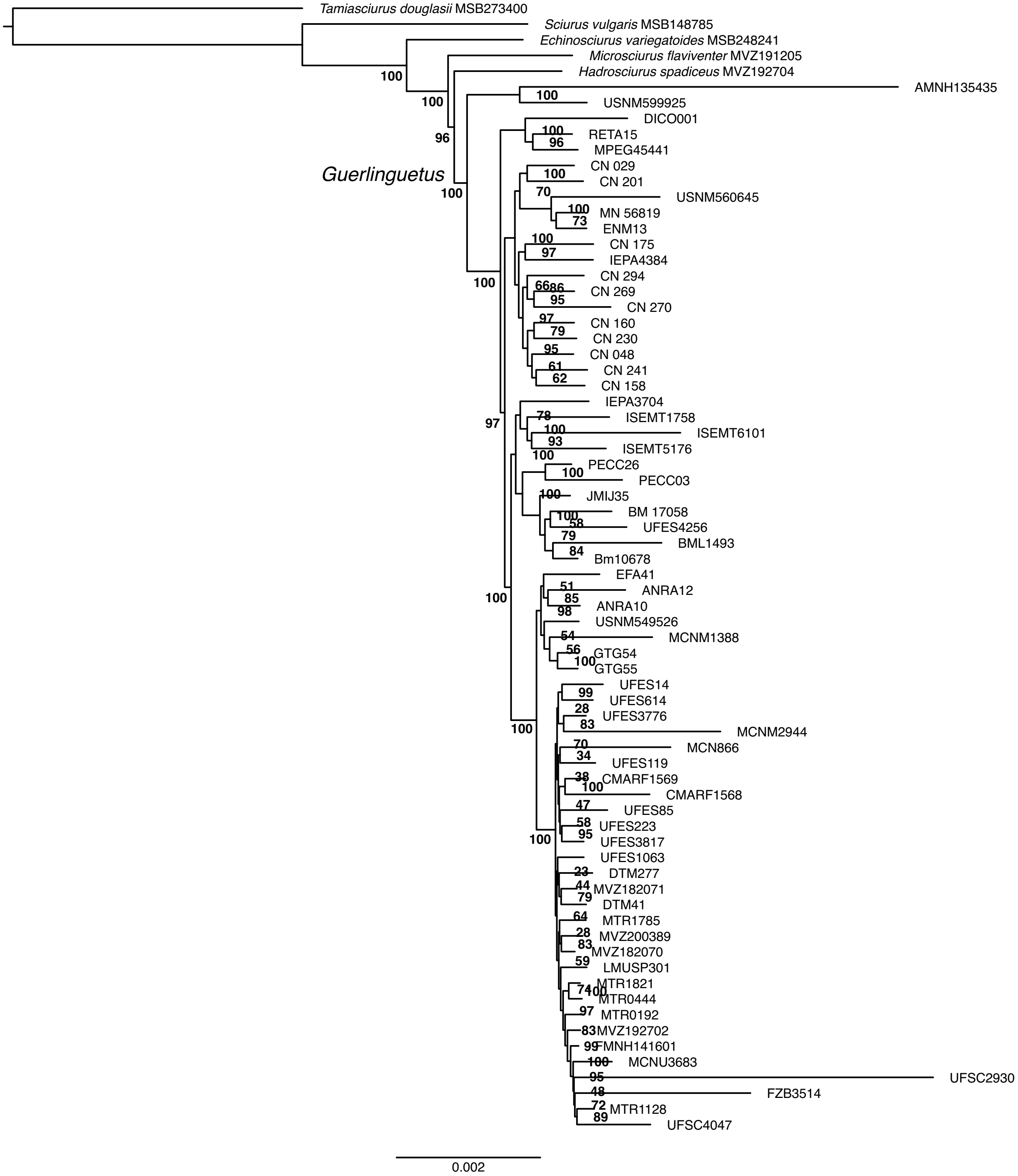

**Supplementary Figure S10**

Phylogenetic reconstruction based on loci (data set including 3,642 UCEs and 67 individuals of *Guerlinguetus*) inferred with SVDquartets.

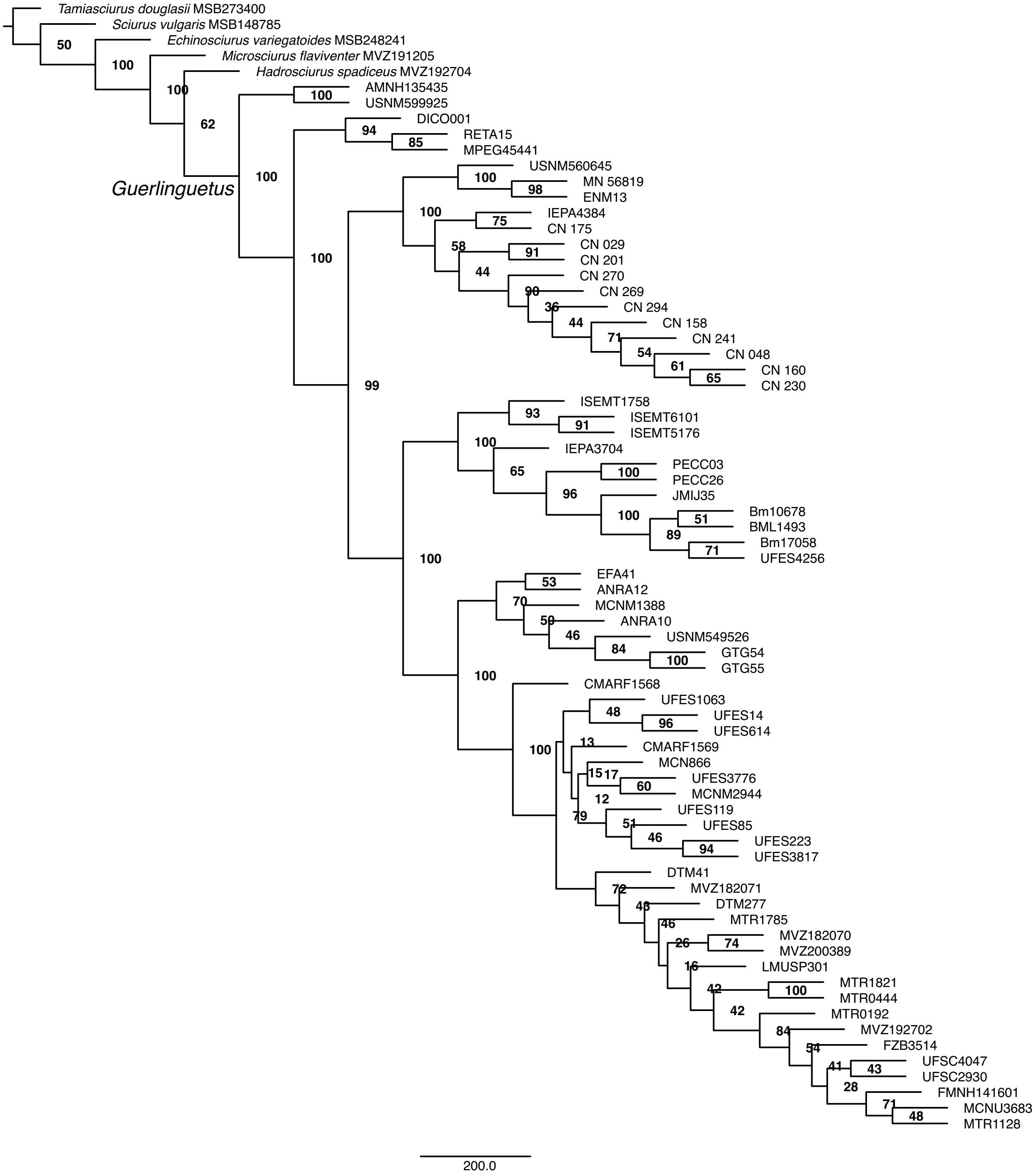

**Supplementary Figure S11**

Model-selection for the TreeMix analysis based on the Δ*m* values obtained for six different models tested (0–5 migration edges). The model including two migration edges better explained the variation in our data according to the Evanno method.

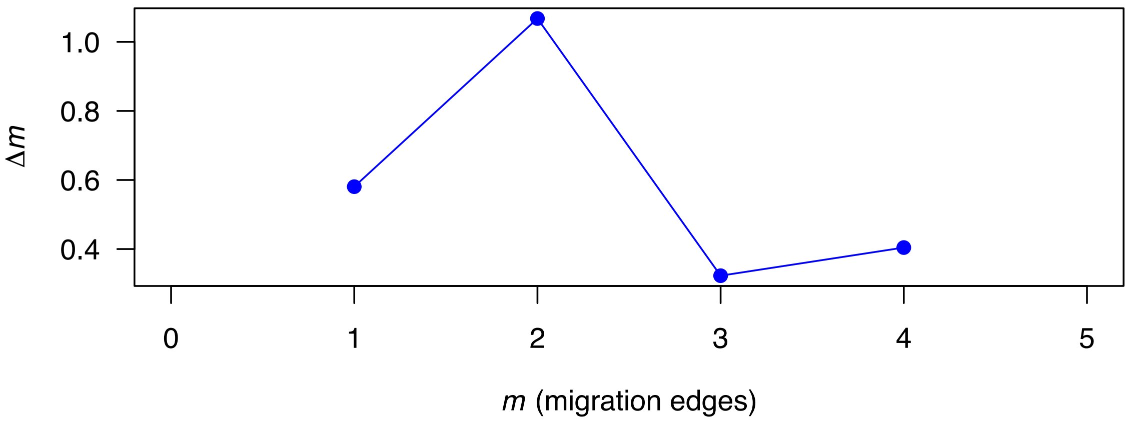
